# Binding of DNA origami to lipids: maximising yield and switching via strand-displacement

**DOI:** 10.1101/2020.06.01.128686

**Authors:** Jasleen Kaur Daljit Singh, Es Darley, Pietro Ridone, James P Gaston, Ali Abbas, Shelley FJ Wickham, Matthew AB Baker

## Abstract

Liposomes are widely used as synthetic analogues of cell membranes and for drug delivery. Lipid-binding DNA nanostructures can modify the shape, porosity and reactivity of liposomes, mediated by cholesterol-modifications. DNA nanostructures can also be designed to switch conformations by DNA strand displacement. However, the optimal conditions to facilitate stable, high-yield DNA-lipid binding while allowing controlled switching by strand-displacement are not known. Here we characterised the effect of cholesterol arrangement, DNA structure, buffer and lipid composition on DNA-lipid binding and strand displacement. We observed that binding was inhibited below pH 4, and above 200 mM NaCl or 40 mM MgCl_2_, was independent of lipid type, and increased with membrane cholesterol content. For simple motifs, binding yield was slightly higher for double-stranded DNA than single-stranded. For larger DNA origami tiles, 4 – 8 cholesterol modifications were optimal, while edge positions and longer spacers increased yield of lipid-binding. Strand displacement achieved controlled removal of DNA tiles from membranes, but was inhibited by overhang domains, which are used to prevent cholesterol aggregation. These findings provide design guidelines for integrating strand-displacement switching with lipid-binding DNA nanostructures. This paves the way for achieving dynamic control of membrane morphology, enabling broader applications in nanomedicine and biophysics.

## INTRODUCTION

DNA nanotechnology is an approach to designing and building nanostructures that self-assemble via DNA hybridisation (1). Since its development (2), a large number of two and three-dimensional DNA nanostructures have been reported (3, 4). The precise addressability of DNA nanostructures allows for patterned functionalisation with nanoparticles (5, 6), proteins (7, 8) and hydrophobic groups (9, 10), resulting in applications in nanofabrication (11, 12), biosensing (13, 14) and membrane targeting (9, 15). Alongside this, in the field of DNA computing, increasingly complex computational circuits have been realised with DNA molecules in solution, driven by the process of toe-hold mediated DNA strand displacement (16, 17). The combination of structural and dynamic DNA nanotechnology has resulted in environment-sensing mechanisms that allow DNA nanostructures to change state in response to external triggers (18).

Modification of DNA with hydrophobic chemical groups, such as cholesterol (9, 19, 20), alkyl chains (21), tocopherol (15), polypropylene oxide (22) and porphyrins (10, 23), has been used to enable lipid membrane binding (24). Cholesterol-modified DNA nanostructures have been used to functionalise liposome surfaces (25), induce membrane curvature and tubulation (26, 27), and form membrane-spanning nanopores that facilitate current flow (9). For example, DNA nanopores can have dimensions which exceed those of natural protein pores (28), and can incorporate mechanisms that regulate ion flow in response to external stimuli, termed gating (20, 29).

A range of lipid-interacting cholesterol-modified DNA nanostructures have been realised to date, but systematic studies of the lipid binding efficiency of cholesterol-modified DNA nanostructures are still incomplete (Figure 1A). The diverse range of design parameters over which to optimise include: DNA nanostructure shape (1D, 2D, 3D) and size (20 - 10,000 bp); hydrophobic modification type, number, position, tether geometry, and strand-displacement reversibility; membrane composition and cholesterol content; buffer components (Na^+^, Mg^2+^); and liposome size and curvature (100 nm – 40 µm). To date, only limited subsets of this parameter space have been explored systematically (Fig. 1A).

**Figure 1:**
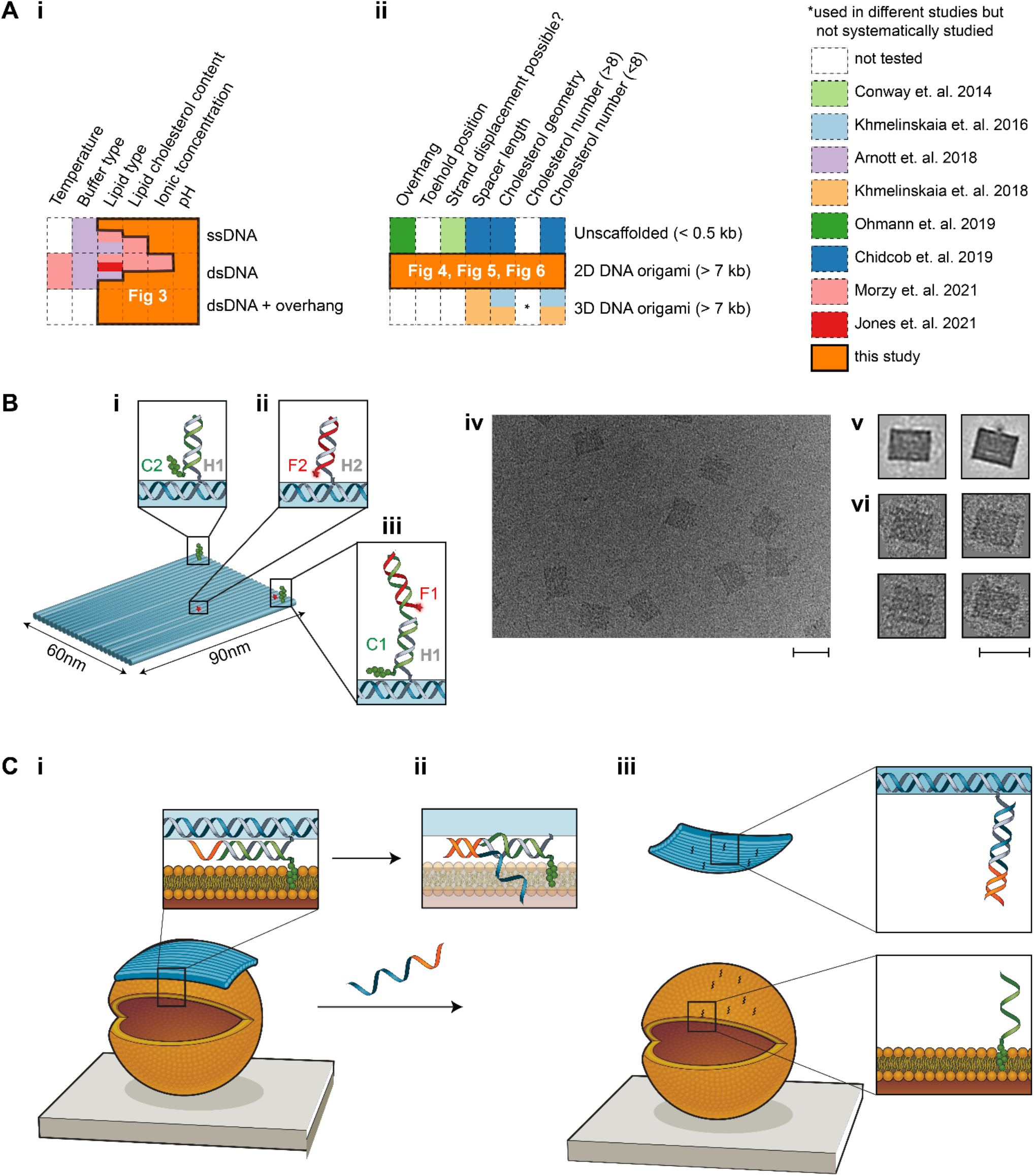
Overview of this work. **(A)** Matrix table highlighting previous studies on membrane binding of cholesterol-modified **(i)** DNA strands and **(ii)** DNA nanostructures, and the gaps filled by our study. **(B)** Schematic of the DNA origami tile. **(i)** Cholesterol (green blobs) labelled strand C2 hybridises with handle H1 on the tile (scaffold strand is shown in blue in insets). **(ii)** Cy5 labelled strand F2 hybridises with handle H2 on the tile. **(iii)** Cholesterol labelled strand C1 hybridises with handle H1 on the tile, as well as Cy5 labelled strand F2. **(iv)** TEM image of the DNA origami tile (no cholesterols). Scale bar: 100 nm. **(v)** Averaged TEM image of tile obtained using RELION. Two different averaged structures were obtained, with minor differences resulting from staining. **(vi)** TEM images of single tiles. Scale bar: 100 nm. **(C) (i)** Schematic of the binding of the tile (blue) to a liposome (orange). Inset: Schematic showing the DNA strand extending from the tile (grey), which is hybridised to a cholesterol strand (green) and has a toehold (orange) for strand displacement. Cholesterol is shown embedded in the lipid bilayer. **(ii)** Upon addition of a displacement strand (blue-orange), branch migration initiates. **(iii)** When strand displacement is complete, the tile is released from the liposome and the cholesterol-DNA remains attached.

The number and position of cholesterol groups on DNA nanostructures has been observed to affect nanostructure docking and diffusion in lipid bilayers (9, 30–32). The effect of cholesterol number and position on membrane binding has been investigated most systemically for a large (∼7000 bp, 110 × 16 × 8 nm) 3D rod-like ‘20 helix-bundle’ nanostructure (30, 31), assembled from a long scaffold strand using the DNA origami folding method (33). It was found that more cholesterol groups increased binding, up to a maximum of 5 tested (30). Cholesterol position was found to be more important than number, with edge-placed groups having the greatest increase in binding. Diffusion on bilayers decreased with the number of corner cholesterol groups. The effect of position and number has also been compared for three DNA origami shapes: a wireframe sphere (80 nm diameter), a long rod (400 × 5 nm), and a flat 2D rectangle (90 × 70 nm) (34). In this case membrane binding was not achieved by hydrophobic-modification of nanostructures, but instead via DNA hybridisation between nanostructure ssDNA ‘overhangs’ and modified ssDNA ‘anchors’ on cell membranes. Similarly, binding increased with overhang number (up to 28) but position was found to be more important, with edge-placed groups providing the largest increase in binding for all shapes. The overall binding yield was determined by comparing nanostructure interaction between cells with and without anchors, and for the 3 shapes the yield was in order of rod, 2D tile, 3D sphere. Small (<500 bp, 7 nm x 7 nm x 7 nm) 3D cube DNA nanostructures, assembled from short synthetic DNA strands (‘unscaffolded’) (32), have also been tested with up to 8 cholesterols. In this case, the mobile fraction of DNA nanostructures on the membrane also decreased with increasing cholesterols, and when cholesterols were on both sides compared to one side.

However, there still remains a gap in the systemic study of cholesterol-modified nanostructures with higher numbers (>8) and different shapes. Many reported membrane interacting nanostructures incorporate much larger numbers of hydrophobic groups, up to 18-26 (9, 15, 28, 35). The large numbers of hydrophobic groups are thought to be necessary to overcome the substantial energy penalties associated with the insertion of membrane-spanning DNA nanopores (36), but have been found to promote aggregation of the modified DNA nanostructures, and reduce yield (15, 37). The shape and size of DNA origami nanostructures have also been shown to affect cellular uptake (38, 39) and membrane binding (30). In particular, the original 2D DNA origami tile (33) is of interest for membrane binding as its modular staple arrangement and large surface area to volume ratio makes it ideal as a molecular pegboard for functionalisation with other molecules, and its flexible geometry (40) gives it the potential to take on the shape and curvature of a membrane.

The effect of the tether length and orientation of hydrophobic groups has also been shown to be important to both binding yield and aggregation of DNA nanostructures. Spacers enhance binding, with a larger increase observed for dsDNA spacers compared to ssDNA, and this effect is greater for designs with fewer cholesterols or centrally positioned cholesterols (31). Unfortunately, spacers also increase the cholesterol-mediated aggregation of DNA nanostructures (41). Adding a ssDNA ‘overhang’ proximal to the cholesterol group has been shown to reduce such aggregation (41). However, the effect of this protective overhang on membrane binding is yet to be quantitatively evaluated. Cholesterol-induced aggregation has also been minimised by labelling cell membranes directly with ssDNA-cholesterol anchors that capture DNA nanostructures from solution (34), but this approach is not suitable for *in vivo* applications. Thus, tether optimisation requires a balance of application type, aggregation, and binding yield, which requires more comprehensive systematic data.

Reversibility in membrane binding allows for complex regulatory mechanisms to be achieved in many biological systems. For instance, amphitropic proteins such as the RAS family (42), hisactophilin (43) and Src kinase (44) utilise reversible membrane binding for regulation of catalytic function and signalling complexes (45, 46). While these systems are present extensively in biology, such reversibility is lacking in DNA-lipid systems. In one study, reversible membrane binding of a wireframe DNA prism nanostructure was achieved using strand displacement (49). The results show that it was necessary to decorate the invader strand with cholesterols for successful displacement of the DNA prism (49). Cation-dependent reversible membrane binding of DNA duplexes has also been demonstrated (50). However, the reversibility of membrane binding is yet to be demonstrated with solid DNA origami nanostructures. Reversible membrane binding could be useful in DNA-assisted liposome formation and purification. Currently, nucleases are standardly used to separate DNA nanostructures from lipid membranes upon liposome formation and purification (47, 48). However, this method may not be suitable for downstream applications that are nuclease sensitive. Reversibility in membrane binding could facilitate nuclease-free removal of DNA nanostructures from liposomes in such instances. DNA toehold design plays an important role in the strand displacement of DNA nanostructures (51, 52), and evidence is emerging that common hydrophobic DNA modifications such as fluorophores can significantly affect strand displacement (53).

Finally, monovalent and divalent cations are necessary buffer components for assembly and stability of DNA duplexes and nanostructures (54, 55), yet are also known to affect the physical characteristics of membrane bilayers (56, 57) and may affect the binding activity of cholesterol-modified DNA (58). Lipid composition plays a further role. For example, for 3D rod nanostructures, using DOPC:DOPS (9:1) decreased membrane binding compared to using DOPC alone (31). For short (<100 bp) DNA oligomers binding to liposomes, ideal ionic conditions have been shown to vary lipid type (50, 58). For PE/PC greater binding was found at 0.3 M KCl, while for PE/PG greater binding was seen in phosphate-buffered saline (PBS), and for both lipid types binding was greater in KCl for dsDNA in comparison with ssDNA (58). The type of hydrophobic modification has also been shown to affect membrane binding. Cholesterol-modified dsDNA demonstrated greater membrane binding when compared with alkyl-phosphorothioate modified dsDNA, on liposomes made from POPC and POPC:DPPC (1:1) (59). Membrane cholesterol content also affected binding with greatest binding for a POPC:cholesterol ratio of 1:1 (vs no cholesterol or a ratio of 1:2).

Membrane binding has also been observed in the absence of cholesterol labelling, with DNA duplexes binding to gel phase DPPC liposomes in the presence of divalent cations such as magnesium and calcium (50). No membrane binding was observed in the presence of monovalent cations alone or in liquid phase membranes (achieved with an increase in temperature or an increase of cholesterol content in the membrane from 0% to 25% or the use of POPC liposomes). Liquid phase POPC membrane binding was possible with the use of cholesterol-labelled ssDNA and dsDNA, with binding increasing as the concentration of magnesium ions increased from 0 mM to 4 mM. Thus, achieving the best match between DNA structure and lipid/buffer conditions remains a challenge.

In this work we fill gaps in our understanding of how DNA-membrane binding is affected by the parameters discussed above. We present a systematic optimisation of the number, position and geometry of cholesterol attachment sites on a 2D rectangular DNA nanostructure, as well as buffer and lipid composition, to improve the efficiency of membrane binding of DNA strands. We have quantified the binding of cholesterol-modified DNA strands to synthetic liposomes using fluorescence microscopy, including the effects of pH, ion concentration, membrane composition and cholesterol content. We investigated three types of DNA motif: a single-stranded DNA (ssDNA), double-stranded duplex (dsDNA), and a duplex with a short single-stranded DNA ‘overhang’ proximal to the cholesterol group, recently shown to reduce aggregation during nanostructure assembly (41). Next, we investigated the membrane binding efficiency of a cholesterol-modified 2D DNA origami nanostructure using fluorescence microscopy and a high throughput gel-shift assay. The effect of cholesterol number, configuration and spacer distance between the DNA nanostructure and cholesterol, were tested. We then optimised strategies for achieving reversible membrane binding by controlled removal of membrane-bound DNA nanostructures using toehold-mediated strand displacement.

## MATERIALS AND METHODS

### Preparation of Buffers and solutions

Liposomes and DNA stocks were diluted in Liposome Buffer (210 mM D-Sorbitol, 5 mM Tris-HCl, pH 7.5) containing NaCl (12.5 mM to 400 mM) and MgCl_2_ (0 mM to 80 mM) as required. For NaCl-dependent experiments, the MgCl_2_ concentration was kept at 10 mM, while for MgCl_2_, the NaCl concentration was kept constant at 100 mM. For pH-dependent experiments a modified Liposome Buffer (210 mM D-Sorbitol, 100 mM NaCl) was used, with pH adjusted to 2, 4, 6, 7, 8 and 10 +/-0.2 with 200 mM NaOH or HCl.

### Design and assembly of ssDNA and dsDNA

DNA strands used for lipid-binding experiments were 23 nucleotides (nt) in length, and used as ssDNA, dsDNA, or dsDNA with a 5’ 6-nt single stranded ‘overhang’ (dsDNA-6nt) (Supplementary Material). DNA sequences were designed using NUPACK design software (60) to prevent unwanted secondary structures. A previously published 6-nt overhang sequence was added to the 5’ end of oligos (41). Oligos were purchased modified at the 3’ end with a tetraethylene glycol cholesterol moiety (TEG-cholesterol), and/or at the 5’ end with Alexa 647, Cy5 or Cy3 fluorophores.

DNA stocks (100 μM, 1000x) were prepared using MilliQ water and stored at 4°C. For dsDNA assembly, non-fluorescent complementary strands were added in a 3-fold excess to fluorescent-modified strands and annealed (90°C for 5 min, then 90-15°C at a rate of -5°C/min) at 10 μM final concentration in duplex buffer (100 mM NaCl, 5 mM Tris-HCl, pH 7.5). Annealed dsDNA were stored at 4°C. DNA was diluted in extrusion buffer (210 mM sorbitol, 100 mM NaCl, 5 mM Tris-HCl, pH 7.5) to 100 nM for lipid-binding experiments. Excess strands were not removed in order to ensure the overall concentration of fluorescent strands remained constant at 100 nM for all samples in all experiments.

### Preparation of Liposomes

Liposomes were produced with two lipid mixtures: (1) DOPE/DOPC liposomes [49.9% 1,2-dioleoyl-sn-glycero-3-phosphoethanolamine, 49.9% 1, 2-dioleoyl-sn-glycero-3-phosphocholin], (2) DPhPC liposomes [99.8% 1,2-diphytanoyl-sn-glycero-3-phosphocholine] (Supplementary Tables 1/2). Both lipid mixtures were doped with 0.1% PE-rhodamine [1,2-dioleoyl-sn-glycero-3-phosphoethanolamine-N-lissamine rhodamine B sulfonyl] for fluorescence imaging and 0.1% PE-biotin [1,2-dioleoyl-sn-glycero-3-phosphoethanolamine-N-biotinyl] for surface tethering. All percentages indicate weight to weight ratios. Liposomes with cholesterol were prepared by replacing either DPhPC (lipid type 2, above) or equal parts of DOPE and DOPC (lipid type 1, above) with cholesterol. All lipids stocks were dissolved in chloroform at 10 mg/mL and stored at -20 °C.

Extruded liposomes, termed small unilamellar vesicles (SUVs) were produced using a Mini-Extruder kit using 100 nm membrane pore size (Avanti Polar Lipids Inc., USA) in matched buffer conditions to the final test conditions (ie, osmolarity and buffer composition matched across the bilayer) and according to the manufacturer’s protocol (Supplementary Methods). Liposomes were then diluted 100-fold to 0.1 mg/mL final lipid concentration in the same buffer in which liposomes were formed prior to experiments (i.e., for 100 mM NaCl, pH 7.5 experiments, extrusion buffer was 210 mM sorbitol, 100 mM NaCl, 5 mM Tris-HCl, pH 7.5).

Giant unilamellar liposomes (GUVs) were prepared by electroformation using the Vesicle Prep Pro machine (Nanion Technologies GmbH, Germany) using the default protocol as described previously (61) (Supplementary Methods). GUVs in electroformation solution (210 mM sorbitol, pH 7.5) were diluted 1:1 in buffer of 210 mM sorbitol, 80 mM NaCl, 10 mM Tris-HCl, giving a final external solution of 210 mM sorbitol, 40 mM NaCl, 5 mM Tris-HCl. Liposome dissolution was tested by titration of increasing concentration of the detergent Polysorbate-20 (Supplementary Figure 1).

### TIRF Fluorescence microscopy of DNA binding to extruded SUVs

Surfaces for imaging were prepared using tunnel slides and BSA-biotin/avidin conjugation chemistry as described previously (62, 63). Briefly, first BSA-biotin was flowed into the tunnel slide to coat and both block the surface and provide sparse, available biotin groups on the surface. Then streptavidin was flowed into the tunnel slide in excess to conjugate to the available biotin groups on the surface. The unbound streptavidin was then washed out and the biotinylated liposomes were flowed in, whereupon the liposome could tether to the available streptavidin binding sites, conjugating the liposome to the surface via BSA-biotin-avidin linkage.

DNA motifs were flowed into the slide upon SUV-tethering on the slide. For DNA-SUV binding experiments, ssDNA or dsDNA strands were flowed into the slide at 100 nM. For experiments with DNA origami tiles, the tiles were flowed in at 10 nM. In both cases, DNA strands or DNA tiles were incubated on the slide for one hour prior to imaging.

Surface-tethered SUVs and DNA were imaged using on a Zeiss Elyra PALM/SIM Microscope in Total Internal Reflection Fluorescence (TIRF) mode with a 63x/1.4 Oil Iris M27 oil immersion objective (Carl Zeiss AG, Germany) and Andor iXon 897 EMCCD camera (Oxford Instruments, United Kingdom). Two-channel fluorescence images were collected for rhodamine-liposomes (excitation/emission filters 561/570-650 plus 750 nm longpass) and fluorophore-tagged DNA (excitation/emission filters 642/655 nm long pass). Exposure times were 100 ms (lipid) and 33ms (DNA), and images were averaged over two subsequent acquisitions.

### Confocal fluorescence microscopy of DNA binding to electroformed GUVs

For DNA-GUV binding, 5 nM of DNA strands were used. DNA binding on GUVs was imaged using a Leica TCS SP8 DLS confocal microscope with HC PL APO CS2 63 x oil immersion objective lens, Acousto-Optical Beam Splitter, and programmable crystal-based beam splitter (Leica Microsystems GmbH, Germany). Two-channel images with rhodamine-liposomes (excitation/emission 561/569-611 nm), and Alexa647-DNA (excitation/emission 640/690-734 nm) and with line averaging of two.

### Quantification of DNA-liposome binding from microscope images

A custom macro script was developed using FIJI in ImageJ (64) to quantify the colocalisation of DNA and liposomes, based on the Manders Overlap Coefficient (65). An intensity threshold was chosen as 2 standard deviations above the mean pixel intensity in the liposome channel over all images from a single experimental condition. This threshold was then used to create a binary mask for assigning pixels as either liposome or non-liposome, and thus identify liposomes from the background (Supplementary Methods, Supplementary Figure 2/3). This method was found to show no bias or correlation with liposome area (percentage coverage), in comparison with Pearson’s correlation, which did show such bias (Supplementary Figure 4/5). Our measurements are ratiometric between liposome-bound and background dye, and we also confirmed that underlying raw fluorescence intensity was not affected by salt concentration (Supplementary Figure 6).

The mean pixel intensities of the DNA channel for the liposome and background areas were then compared, and used to calculate a colocalisation ratio, C_R_, via:

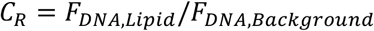

Where *C*_R_ is the reported colocalisation ratio, F_DNA,Lipid_ is the mean pixel intensity of the fluorescent DNA in the liposome region of the DNA channel, and F_DNA,background_ is the mean pixel intensity of the fluorescent DNA in the background region of the DNA channel (Supplementary Figure 3).

### Assembly of DNA origami nanostructures

To decorate the DNA tile with cholesterols, 21-nt ssDNA handles (H1) were designed for hybridization with complementary cholesterol-TEG modified DNA strands (C1 or C2). To quantify binding of cholesterol-DNA to the tile, strand C1 was designed with a second binding domain for hybridisation of Cy5 fluorophore labelled DNA (F1) (Figure 1B-iii). For membrane binding experiments, strand C2, without the second domain, was used (Figure 1B-i). To label the tile with fluorophores for microscopy, separate 21-nt ssDNA handles, H2 were extended from the surface of the tile for hybridisation with Cy5 labelled ssDNA F2 (Figure 1B-ii). In this design for microscopy, the number of fluorophores on the tile is independent of the number of cholesterols. The positions of H1 on the tile are given in Figure 4A and Supplementary Figure 9, and the positions of H2 are given in Supplementary Figure 10.

DNA sequences for DNA origami tile structure (33) were obtained using the Picasso software (66). The tile was folded using 10 nM of M13mp18 ssDNA scaffold and 10x excess (100 nM) of DNA staple strands in folding buffer (5 mM Tris, 1 mM EDTA, 12 mM MgCl_2_, pH 8.0) and annealed over 3 hours (80°C for 15 min, then 60– 4°C in 56 steps at 3 min 12 sec/step). Cholesterol-TEG modified DNA strands (C1 or C2) were added to staple pools prior to annealing at 2x excess relative to staple concentration (e.g. for 4C tile, C1/C2 = 100 nM x 4 × 2 = 800 nM). For fluorophore attachment to cholesterol strands, F1 strand was added at 2x excess to the amount of C1 (e.g. for 4C tile, F1 = 800 nM x 2 = 1600 nM). For fluorescence labelling of tile, 2000 nM of F2 strand was added as there are ten H2 handles on the tile (so, F2 = 100 nM x 10 × 2 = 2000 nM). All staples, plain and modified, were annealed in one-pot. All sequences used are supplied in the Supplementary Data.

### Purification of DNA origami by agarose gel electrophoresis

For analysis of membrane binding by gel, DNA origami tiles were purified by agarose gel electrophoresis (67). Samples were loaded on 2% agarose gels and run for 2.5 hours at 60 V at RT. Gel and running buffer used was 0.5× TBE buffer (45 mM Tris boric acid, 45 mM Tris base, 1 mM EDTA, pH8) with 11 mM MgCl_2_, gels were pre-stained with SyBrSafe stain. Gels were viewed under an LED Blue Light Transilluminator (Fisher Biotec) and the bands corresponding to the DNA origami tile were cut. The cut bands were transferred into Freeze ‘N Squeeze DNA Gel Extraction spin columns (Bio-Rad), crushed and extracted by centrifugation at 18,000g and 4°C for 10 minutes. The concentration of the recovered solution was determined using a Nanodrop (Thermo Fisher Scientific) to measure absorption at 260 nm. DNA origami were stored at 4°C.

### Purification of DNA origami by PEG precipitation

DNA origami tiles were purified by PEG precipitation (68) for all microscopy experiments. The folded DNA origami tile sample was mixed at 1:1 ratio with PEG buffer (15% PEG 8000 (w/v), 5 mM Tris, 1 mm EDTA, and 505 mM NaCl) and incubated at 4°C for 30 minutes. The solution was centrifuged at 15,000g at 4°C for 30 minutes. The supernatant was removed using a pipette and discarded. The remaining pellet was air dried and then dissolved in the buffer (5 mM Tris-HCl, 40 mM NaCl, 10 mM MgCl2).

### Gel-shift assay to quantify membrane binding of DNA origami

Agarose gel shift assays were conducted to determine the extent of membrane binding of DNA origami nanostructures (58, 69). 10 µL of 2.5 nM gel purified DNA origami tile was incubated with 5 µL of extruded SUVs (diluted 20x in 5 mM Tris-HCl, 40 mM NaCl, 10mM MgCl_2_ upon extrusion) for 30 minutes at room temperature. 10 µL of sample was loaded onto a 2% agarose gel prepared in 0.5× TBE buffer (45 mM Tris boric acid, 45 mM Tris base, 1 mM EDTA, pH 8.0) supplemented with 11 mM MgCl_2_. The gel was run at 60 V for 2.5 hours at 20°C, and imaged using Chemidoc MP Imager (Bio-rad). Images were obtained in the Cy5 channel and analysed using the Bio-Rad Image Lab software.

This assay allowed for the separation of unbound tiles from membrane bound tiles (Figure 4C). For each tile design, 2 gel lanes were run: (1) tiles incubated with liposomes (sample lane), (2) tiles only, no liposomes (control lane). The intensity of the tile band was compared between the sample and control lanes to quantify the extent of membrane binding. The ratio of the intensity of the tile band in the sample lane (+ liposomes, red box in Figure 4C.i.) to the intensity of the tile band in the control lane (-liposomes, black box in Figure 4C.i) was determined to calculate the percentage of bound tiles:

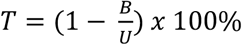

where T is the estimated percentage of tiles that are bound to the membrane, B is the intensity of the tile band in the lane with liposomes, and U is the intensity of the tile band in the lane without liposomes. To account for experimental variation in loading of DNA into the gel, at least two repeats of each gel were conducted.

### Transmission electron microscopy of DNA origami

15 µL of 1 nM (in 5 mM Tris, 1 mM EDTA, 12 mM MgCl_2_, pH 8.0) purified DNA origami tile sample was placed onto a parafilm, a plasma treated carbon-coated TEM grid (Ted Pella EM grids from ProScitech) was placed onto the sample and left for 1 min for the sample to adsorb onto the grid. A 2 µL droplet of 2% uranyl acetate solution was placed onto a fresh parafilm and the grid was then quickly tapped onto the droplet and immediately tapped onto a filter paper to remove excess stain. This staining protocol was repeated three times. TEM imaging was performed using the JEOL JEM-1400 microscope, 120 kV. TEM micrographs of the DNA tiles were averaged using RELION (70) (Supplementary Figure 11).

## RESULTS

### DNA origami nanostructure design and assembly

The 2D rectangle DNA origami tile (Fig. 1B) was chosen due to its ease of assembly and wide use in the field of DNA nanotechnology (37, 66, 71). The tile consists of 24 parallel DNA helices folded using the M-13 scaffold and has dimensions of 60 nm x 90 nm x 2 nm. Successful assembly of the tile and incorporation of fluorophore-modified staple strands was verified using agarose gel electrophoresis (Supplementary Figure 8) and transmission electron microscopy (TEM) (Fig. 1B-iv and Supplementary Figure 11).

### Imaging of DNA-liposome binding

Sample homogeneity for SUVs was characterised using light scattering (Supplementary Figure 7) and then colocalisation of fluorescent DNA to fluorescent liposomes was measured using TIRF microscopy. Colocalisation was observed only when DNA oligomers were modified with cholesterol (Fig. 2). In contrast, DNA without cholesterol was distributed evenly throughout the image independently of the position of liposomes (Fig. 2A). Similarly, in confocal images of GUVs, plain DNA did not colocalise (Fig. 2B) whereas cholesterol-modified DNA colocalised with the GUVs (Fig. 2D).

**Figure 2:**
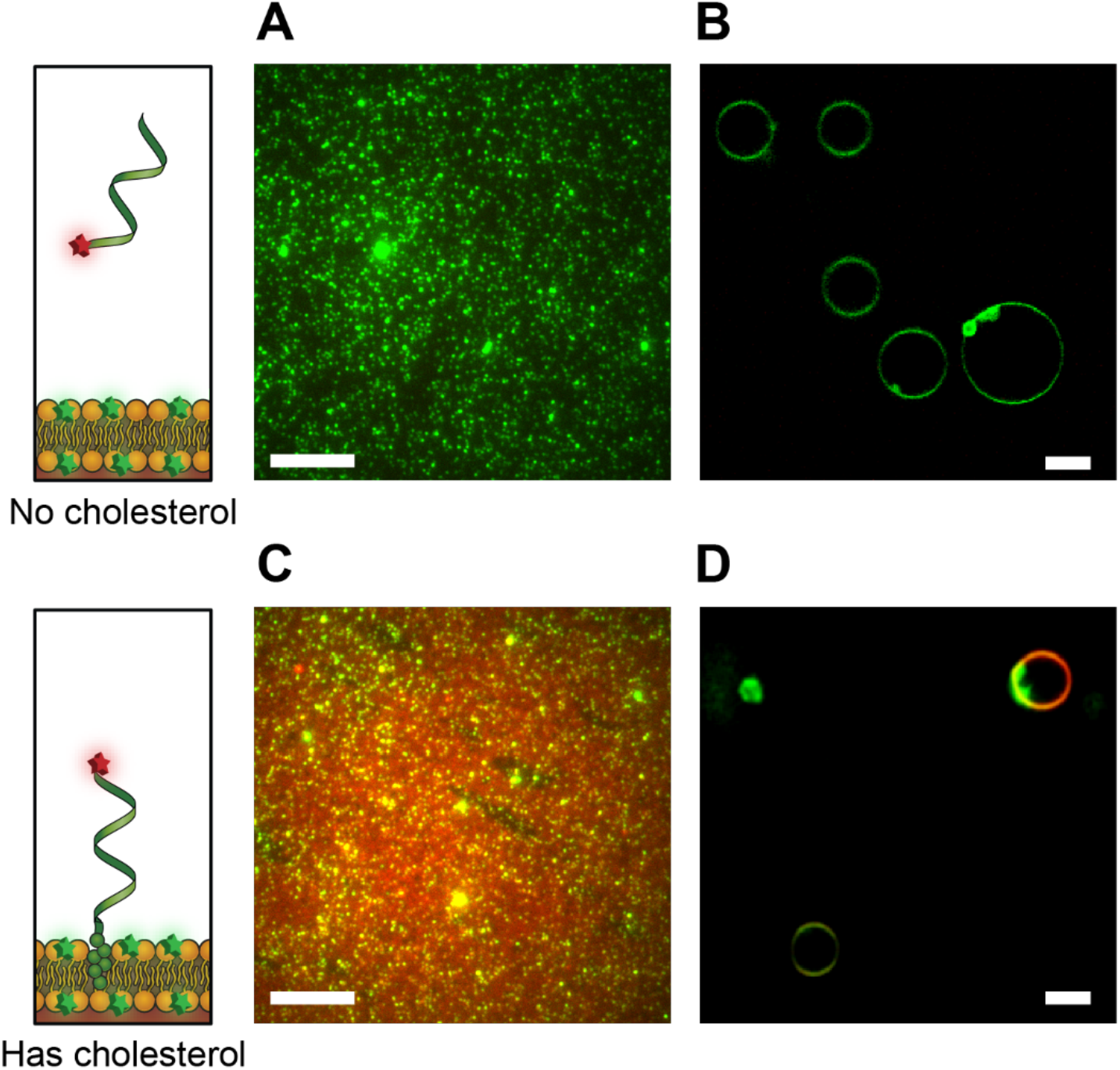
Binding of cholesterolated DNA to liposomes. **(A)** Merged two-colour TIRF images of ssDNA Alexa647-DNA oligomer (red) at 100 nM concentration and PE-rhodamine-labelled DOPE/DOPC SUVs (green). C_R_ for image in (A) is 1.0332. **(B)** Confocal image of ssDNA Alexa647-DNA oligomer (red) at 5 nM and rhodamine-labelled DOPE/DOPC GUVs (green). **(C)** Merged two-colour TIRF images of ssDNA Alexa647-DNA-cholesterol oligomer (red) at 100 nM concentration and PE-rhodamine-labelled DOPE/DOPC SUVs (green). C_R_ for image in (C) is 1.9436. **(D)** Confocal image of ssDNA Alexa647-DNA-cholesterol oligomer (red) at 5 nM and rhodamine-labelled DOPE/DOPC GUVs (green). For TIRF images (A/C, 16 bit) brightness are scaled with min = 1500, max = 7000 au. For GUVs image scaled (B/D, 8 bit) min = 30, max = 255. For all images, buffer conditions are 210 mM sorbitol, 5 mM Tris-HCl, pH 7.5; for A/C NaCl 25 mM, for B/D NaCl 50 mM. All scale bars: 10 μm.

### Effect of buffer and lipid composition on DNA-liposome binding

The effect of DNA origami buffer components on the binding of cholesterol-DNA to SUVs was quantified. The colocalisation ratios (C_R_) of cholesterol-modified ssDNA, dsDNA and dsDNA-6 nt were measured for varying [NaCl], [MgCl2] and pH, for 1:1 DOPE/DOPC liposomes and DPhPC liposomes, and compared to plain DNA controls (Fig. 3).

**Figure 3:**
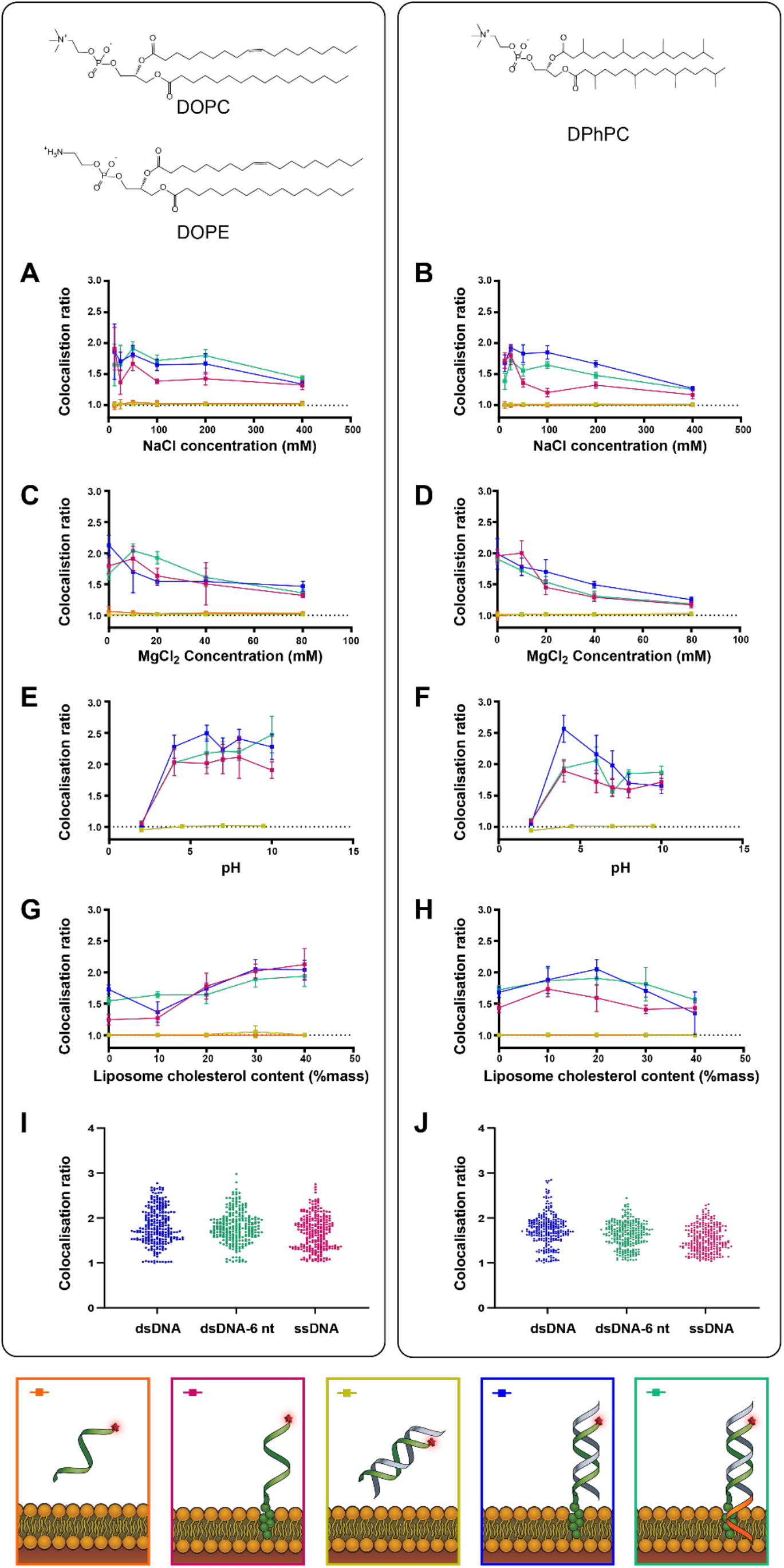
The effect of NaCl, MgCl_2_, pH, and membrane cholesterol on DNA-liposome colocalization in SUVs. Colocalisation ratios and standard deviations are shown for Alexa647-labelled cholesterol-tagged single stranded DNA (ssDNA, pink), cholesterol-tagged double stranded DNA (dsDNA, blue) and cholesterol-tagged double stranded DNA with a 6 nt overhang (dsDNA-6nt, green) as well as dsDNA with no cholesterol tag (yellow) and ssDNA with no cholesterol tag (orange) and rhodamine-labelled DOPE/DOPC liposomes **(left column, A/C/E/G)** and DPhPC liposomes **(right column, B/D/F/H)**. Solution conditions tested included: **(A/B)** extrusion buffer [NaCl] containing 12.5, 25, 50, 100, 200 and 400 mM NaCl, **(C/D)** extrusion buffer [MgCl2] containing 0, 10, 20, 40 and 80 mM MgCl2, and **(E/F)** extrusion buffer [pH] adjusted to pH values of 2, 4, 6, 7, 8 and 10. **(G/H)** The effect of lipid cholesterol content was tested by forming liposomes from DOPE/DOPC and DPhPC lipid stocks containing 0, 10, 20, 30 or 40% cholesterol. **(I/J)** Distribution of C_R_ values for each cholesterol-tagged DNA configuration across all conditions (n = 264; DOPE/DOPC: dsDNA = 1.82 ± 0.41; dsDNA-6nt = 1.80 ± 0.35; ssDNA 1.68 ± 0.37; DPhPC: dsDNA = 1.74 ± 0.36, dsDNA-6nt = 1.63 ± 0.28, ssDNA = 1.53 ± 0.29, all mean ± SD). For both lipid types and with or without overhang, ssDNA vs dsDNA showed a significant difference in mean (Wilcoxon rank sum test, p<0.05). For DPhPC only there was a significant difference in mean, albeit smaller than the associated error measurement (△C_R_ = 0.10 ± 0.46) between dsDNA and dsDNA-6nt (Wilcoxon rank sum test, p<0.05).

DNA that was evenly distributed throughout a slide independently of liposome location would be expected to produce a C_R_ value of 1.0. Membrane-bound DNA, on the other hand, would be expected to produce a C_R_ of greater than 1.0, as the concentration of DNA on liposomes is expected to be higher than the concentration on the background. For DNA without cholesterol, the C_R_ was measured as 1.01 ± 0.03 for all NaCl, pH, MgCl_2_ and lipid conditions tested, indicating low non-specific interaction between the DNA and liposomes. In each case, we compared colocalization ratio of cholesterolated dye-DNA with dye-only DNA demonstrating that the non-specific background binding from the dye was negligible. In comparison, the colocalisation ratio for cholesterol-DNA configurations when pooled across all conditions was significantly higher than without cholesterol (C_R_ = 1.70 ± 0.36, p<0.05; excluding measurements for cholesterolated DNA at pH = 2.0 measurements yields C_R_ = 1.73 ± 0.34, p<0.05). This confirmed that specific binding of cholesterol-DNA to liposomes was observed.

For all three configurations of cholesterol-DNA (ssDNA, dsDNA, dsNDA-6nt) on both DOPE/DOPC liposomes and DPhPC liposomes, a significant decrease was observed in C_R_ between 12.5 mM and 400 mM NaCl and between 0 mM and 80 mM MgCl_2_. Linear regression analysis for all three configurations on both liposome compositions showed a trend of decreasing colocalisation scores with increasing concentrations of NaCl and MgCl_2_ (95% CI of gradient < 0).

DNA-liposome binding was tested for pH values between 2 and 10 and the C_R_ of cholesterol-DNA was observed to decrease in highly acidic conditions. At pH 2, the C_R_ of all three configurations of cholesterol-DNA with both DOPE/DOPC liposomes (Fig. 3E) and DPhPC liposomes (Fig. 3F) decreased to a level similar to the non-cholesterol control strands, and was significantly less than at all other pH values (p < 0.05). This indicates that solutions of pH 2 inhibit membrane binding.

### Effect of DNA configuration on lipid binding

Significant differences were observed in mean C_R_ for the different DNA configurations. In both DOPE/DOPC liposomes and DPhPC liposomes, we found dsDNA to colocalise more than ssDNA, in the order C_R(dsDNA)_ ≈ C_R(dsDNA-6 nt)_ > C_R(ssDNA)._ When compared to dsDNA, the addition of the six nucleotide overhang in dsDNA-6 nt was observed to cause a modest but significant decrease in binding to DPhPC liposomes, but no significant decrease in binding to DOPE/DOPC liposomes (Fig. 3I-J).

### Effect of membrane cholesterol content on DNA-lipid binding

Cholesterol content was increased between 0% and 40% by mass for both lipid compositions. For DOPE/DOPC liposomes, C_R_ of all three configurations of cholesterol-DNA showed a significant increase between 0% and 40% cholesterol (p < 0.05). Linear regression analysis showed a trend of increasing C_R_ across the observed range of membrane cholesterol content (gradient 95% CI > 0) (Fig. 3G).

For DPhPC liposomes, C_R_ of cholesterol-tagged DNA increased to a maximum at 10%-20% membrane cholesterol, then decreased with further increasing cholesterol. All three configurations of cholesterol-DNA showed both a significant increase in C_R_ between 0% and 20% (p < 0.05) and a significant decrease in C_R_ between 20% and 40% (p < 0.05) (Fig. 3H).

### Effect of number of cholesterols on membrane binding of DNA origami tile

First, the correct assembly of cholesterol-DNA (C1) (Fig. 1B) to the DNA tile was verified using agarose gel analysis (AGE) (Supplementary Figure 12) and TEM (Supplementary Figure 13). The tile was folded with either 0, 1, 2, 4, 8 or 16 cholesterols (0C, 1C, 4C-LS, 8C, 16C) moieties (Fig. 4A and Supplementary Figure 9). Cy5 fluorophore-labelled strand F1 is designed to hybridise to the tile only in the presence of cholesterol-DNA C1. A comparison was made of Cy5 intensity of the DNA tile band (normalised to SyBr safe stained DNA channel) for samples with different numbers of cholesterol-DNA sites. The normalised Cy5 intensity was found to increase with increasing cholesterol number (Supplementary Figure 12). This successfully confirmed that more cholesterol-DNA strands attached to the tile as the number of handles (H1) increased from 0 to 16.

**Figure 4:**
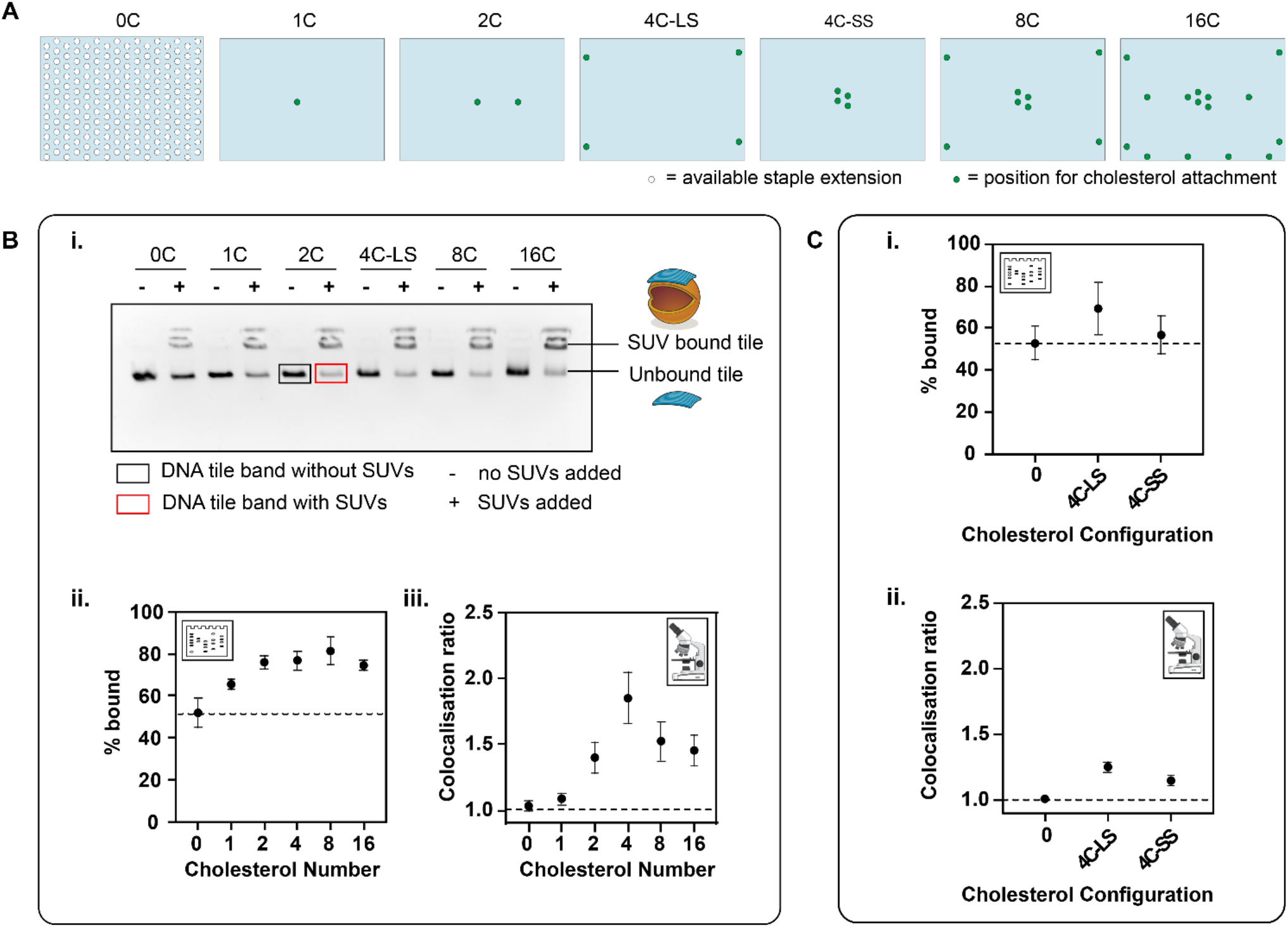
Effect of number and arrangement of cholesterols on membrane binding. **(A)** The different attachment points available on the DNA origami tile (light blue rectangle) for staple extensions (white circle) and the points selected for staple extension of handles H1 (green circle) for cholesterol attachment. **(B)** The effect of cholesterol number on membrane binding. **(i)** Gel image. **(ii)** Percentage bound from the gel analysis calculated from ratio of integrated band intensity in the presence of liposomes (red box) to the integrated band intensity in the absence of liposomes (black box). **(iii)** Colocalisation ratios from microscopy. **(D)** The effect of cholesterol configuration on membrane binding. **(i)** Percentage bound obtained from gel analysis. **(ii)** Colocalisation ratios from microscopy.

The effect of cholesterol number on membrane binding was then observed by AGE and fluorescence microscopy (TIRF). In AGE, the DNA tiles were observed to migrate through the gel matrix, while the liposomes remained in the wells (Supplementary Figure 14), in agreement with literature results using this technique (41). The percentage of tiles bound to the membrane was 52 ± 5% for tile-0C, indicating significant non-specific membrane binding even in the absence of cholesterols (Figure 4B-ii and Supplementary Figure 15). A significant increase in percentage of tiles bound was observed from tile-0C to tile-8C (p<0.01). Maximum binding of tiles to liposomes was observed for tile-8C at 81 ± 5%.

For microscopy experiments, the maximum colocalisation of tile and liposomes was observed on tiles with four cholesterol groups (C_R_ = 1.85 ± 0.20) (Fig. 4B.iii). A significant increase in C_R_ was observed between n = 0 and n = 4 cholesterol groups (p<0.05), with a trend of increasing colocalisation as the number of cholesterol groups was increased within this range (linear regression between n=0 and n=4: gradient 95%CI > 0). For the control with no cholesterol, *C*_R_ = 1, indicating similar density of DNA in the background and on the liposome. Linear regression analysis across n = 4, n = 8 and n = 16 showed a decreasing trend as the number of cholesterol groups was increased (gradient 95%CI < 0), however there was no significant difference in the means from pairwise testing.

### Effect of cholesterol geometry on membrane binding of DNA origami tile

We next investigated the effect of cholesterol-DNA geometry on membrane binding. Two different geometries were compared: Large Square (4C-LS) and Small Square (4C-SS), as shown in Fig. 4A-i. In the LS configuration, four cholesterol anchors were positioned along the edge of the tile. The separation between the handles in the 4C-LS configuration is 80 nm along the long edge and 45 nm along the short edge of the tile. In the 4C-SS configuration, four cholesterols anchors were positioned at the centre of the tile, with a separation of 5 nm between the handles. The percentage of tiles bound to the membrane was 69 ± 12 % and 57 ± 9% for 4C-LS and 4C-SS configurations, respectively, using the gel-shift assay (Fig. 4C-i and Supplementary Figure 16). The percentage of tiles bound non-specifically in the no cholesterol sample was 53 ± 8 %, similar to results discussed in the previous section. The differences in membrane binding between the no cholesterol control, the 4C-LS and the 4C-SS geometries were found not to be statistically significant.

For the microscopy assay, C_R_ of 1.25 ± 0.04 and 1.15 ± 0.03 were obtained for the 4C-LS and 4C-SS configuration, respectively (*t*-test p<0.05, Fig. 4C-ii). These C_R_ values are lower than those previously observed for the tile with 4 cholesterols shown in Fig. 4B-iii, but we note that the C_R_ values vary with different SUV preparation batches but are consistent within a single SUV preparation.

The extent of cholesterol attachment for the 4C-LS and 4C-SS geometries were measured by AGE (method as above, Supplementary Figure 17) to determine if the difference observed in membrane binding between these samples was due to different levels of cholesterol attachment to the tile. No differences in cholesterol attachment were observed for the two tile configurations, suggesting that the differences observed in lipid binding were due to the different geometrical arrangements of the cholesterols on the tile.

### Effect of spacer length between cholesterol and tile on membrane binding

Next, the effect of the spacer length between the cholesterol and the tile on membrane binding was observed. Seven different spacer designs were tested: Dt1.4, Dt6.1, Dt8.1, Pt8.1, Pt8.5, Pt13.2, and Dt15.2, where the number gives the distance between the cholesterols and the tile, e.g. Dt15.2 represents an estimated spacing of 15.2 nm. The initial (Dt or Pt) refers to the positioning of the toehold relative to the cholesterol, either distal to the cholesterol (Dt) or proximal to the cholesterol (Pt). Design Dt15.2 has an additional 10-nt overhang next to the cholesterol group, which is expected to reduce aggregation (41).

Membrane binding experiments were performed using the gel shift assay (Fig. 5B-i and Supplementary Figure 18). Of the seven designs tested, the lowest percentage of bound tiles was observed for Dt1.4 at 48% ± 27%. The maximum percentage of bound tiles was observed for Dt15.2 at 89% ± 9%. Percentage of bound tiles for Dt6.1, Dt8.1, Pt8.1, Pt8.5, Pt13.2 was at 50 ± 15%, 62 ± 20%, 67 ± 23%, 80 ± 23% and 86 ± 15%, respectively. The control with no cholesterols had 40 ± 22% tiles bound to the membrane in this experiment. For microscopy, only three designs were selected: Dt1.4, Pt8.5 and Dt15.2. C_R_ of 1.07 ± 0.05, 1.15 ± 0.04 and 1.24 ±0.04 were obtained for Dt1.4, Pt8.5 and Dt15.2, respectively (Fig. 5B-ii). The percentage of tiles bound in the presence and absence of Cy5 fluorophores was compared, and no significant difference was observed (Supplementary Figure 19).

**Figure 5:**
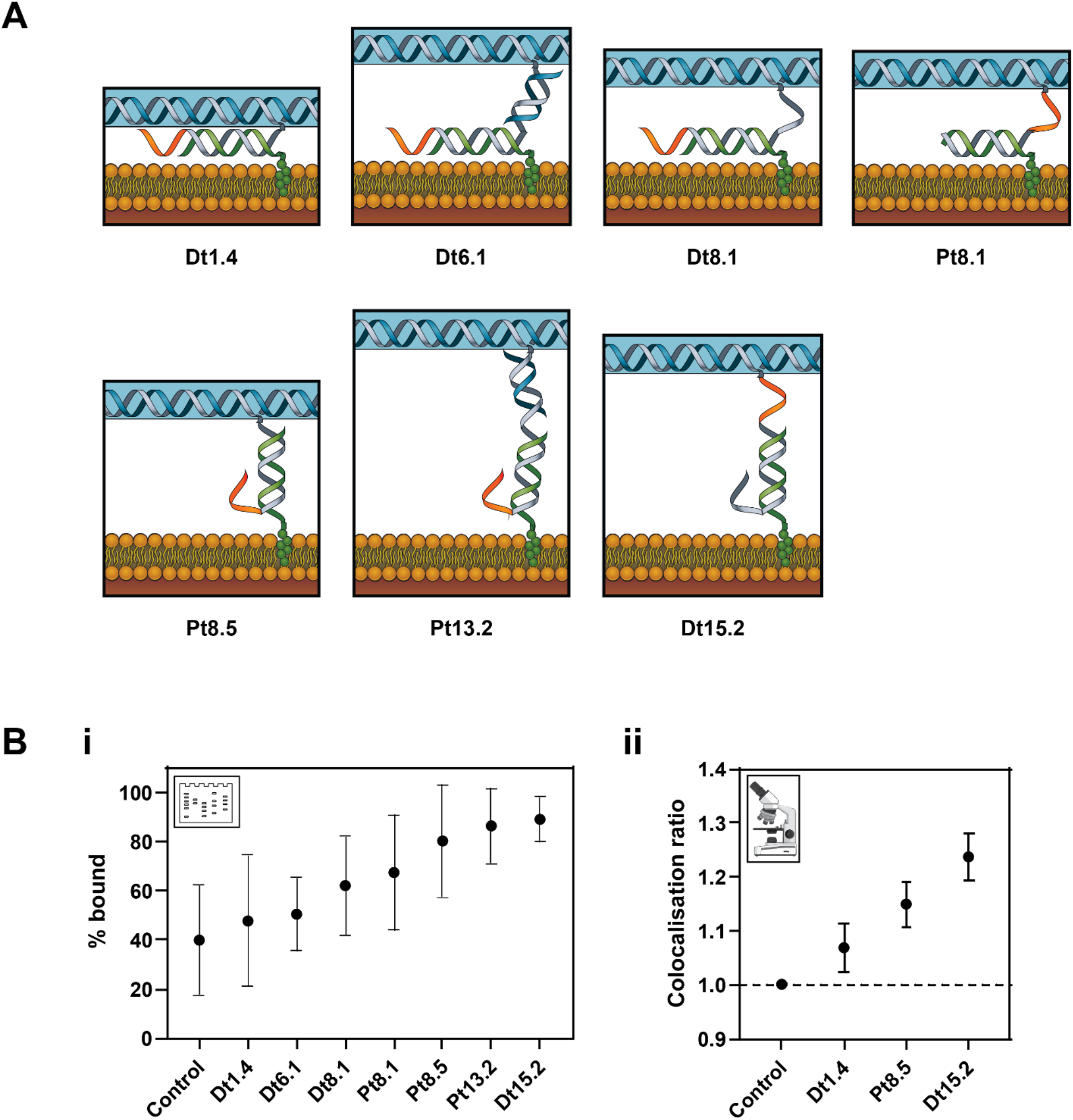
Effect of spacing and linker length between cholesterol and DNA origami tile on binding to liposomes. **(A)** Schematics for the different designs for cholesterol attachment to the tile and its membrane binding. **(B)** The effect of spacing between cholesterol and tile on the membrane binding. **(i)** Percentage bound obtained from gel analysis. **(ii)** Colocalisation ratios from microscopy.

### Effect of toehold position on strand displacement of cholesterol-DNA from DNA origami tiles

The effect of toehold position on releasing DNA tile binding to SUVs by strand displacement was then investigated for the 4C-LS configuration. A 10-nt toehold was used for strand displacement of the tile from SUVs, to facilitate displacement of the C2 cholesterol strand from the tile (Fig. 1C). The toehold was designed to be either proximal (Pt) or distal (Dt) to the cholesterol modification.

Strand displacement was validated by folding DNA tiles with cholesterols and fluorophore (Cy5) such that the fluorophore attachment was dependent on the cholesterol attachment, similar to earlier experiments used to confirm cholesterol attachment (Fig. 1B-iii, Supplementary Figure 12). The strand arrangement for each of the different designs and their toehold positions is shown in Fig. 6A-i.

**Figure 6:**
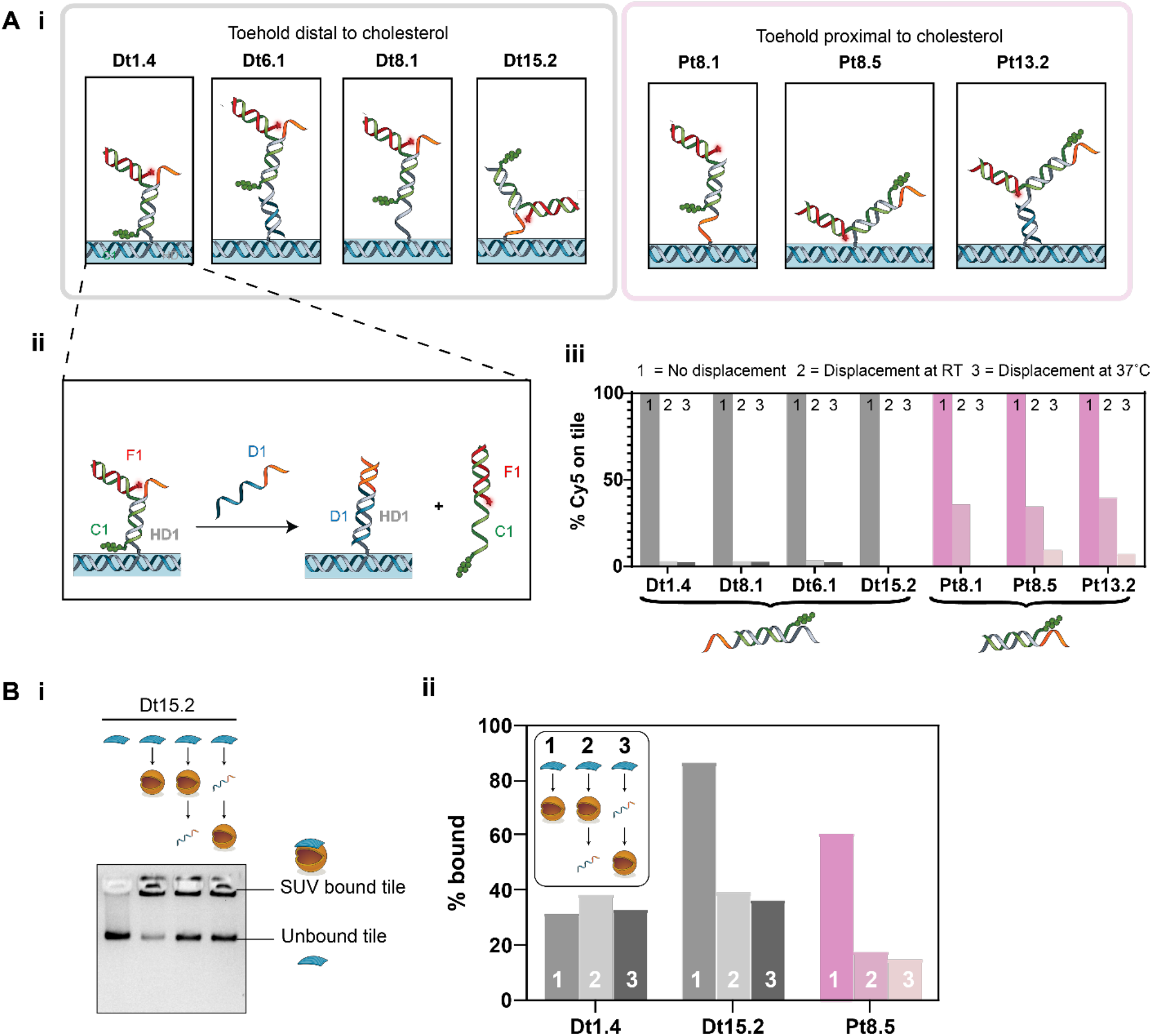
Effect of toehold position on strand displacement release of DNA tiles from liposomes. **(A) (i)** Schematics for the different designs for cholesterol attachment to the tile. **(ii)** Example strand displacement mechanism resulting in the detachment of the Cy5 labelled strand from the tile, shown here for Design Dt1.4. **(iii)** Bar chart showing the percentage of Cy5 attached to the tile under different displacement conditions. In the control, no displacement strand is added, RT and 37°C represent displacement at room temperature and 37°C, respectively. **(B)** Strand displacement of membrane-bound tiles. **(i)** Gel image for Dt15.2. **(ii)** Percentage of tiles bound to the SUVs tabulated from the gel analysis. Designs with toeholds distal from the cholesterol groups are shown in shades of grey. Designs with toeholds proximal to the cholesterol groups are shown in shades of pink.

Toehold-mediated strand displacement of the cholesterol strand from the DNA tile was first validated without any binding to SUVs. When displacement strand D1 is added, it is expected to bind to the toehold on the handle H1 and initiate branch migration, displacing the C1-F1 complex. It was confirmed that strands C1-F1 were displaced from the tile on addition of D1, as indicated by decrease in Cy5 intensity of the DNA tile band in AGE (Fig. 6A-iii, Supplementary Figure 20).

Displacement efficiency was compared at room temperature (RT) and 37°C for 30 mins using 4 µM of strand D1 and 2.5 nM of tile. The percentage of Cy5 on the tile was determined from the gel as shown in Fig. 6A-iii. The percentage of Cy5 on the tile for the no displacement control was taken as 100% and the percentages for the displacement samples were calculated relative to this. Greater displacement was observed for designs with the toehold distal to the cholesterol groups (Fig. 6A-iii and Supplementary Figure 20). For the tested temperatures, we observed no significant change in the displacement of the designs with distal toeholds, which were all highly efficient, decreasing from 100% to <3% for all conditions tested. In contrast, displacement efficiency of designs with toehold proximal to the cholesterol groups showed temperature dependence. Only partial displacement was observed at room temperature (decrease from 100% to 34-39%), with improved displacement at 37°C (100% to 0-9%).

### Effect of toehold position on strand displacement of DNA-tiles from liposomes

Strand displacement of lipid-bound DNA tiles was then tested. For these experiments, cholesterols were attached to the tile (Fig. 1B-i) independently of the fluorophores (Fig. 1B-ii). The effect of strand displacement of the cholesterol from the tile was compared before (pre-displaced) and after (displaced) incubation with liposomes.

A subset of designs previously tested for lipid binding (Fig. 5B) and strand displacement in lipid-free conditions (Fig. 6A-iii) were selected, consisting of Designs Dt1.4, Dt 15.2 and Pt8.5. Previously, Dt1.4 had shown poor membrane binding but efficient strand displacement in lipid-free conditions. Dt15.2 had both high membrane binding and efficient strand displacement. Pt8.5 had high membrane binding but inefficient strand displacement at room temperature.

The percentage of lipid bound tiles for each design was determined by gel analysis for each experimental condition (Fig. 6B and Supplementary Figure 21). The results show Dt1.4 had poor initial membrane binding (32%), which made it difficult to determine if there was a change in binding after strand displacement (38%, displaced, 33%, pre-displaced). Dt15.2 was observed to have both high initial membrane binding (86%) and a decrease in binding on strand displacement indicating successful displacement (39%, displaced, 36%, pre-displaced). Pt8.5 had moderate initial membrane binding (61%) and successful strand displacement (18%, displaced, 15%, pre-displaced). Comparing pre-displaced and displaced values, for all samples no significant effect was observed on changing the order of lipid binding and displacement. Hypothesised interactions of Dt1.4, Dt15.2 and Pt8.5 for both the bound and unbound tiles are proposed in Supplementary Figure 23.

## DISCUSSION

Our comparison of the lipid-binding yield of simple DNA motifs (ssDNA, dsDNA, dsDNA-6nt) and large DNA origami tiles (7249 bp) modified with cholesterol defines a range of experimental conditions for which binding yield is independent of external buffer and lipid composition. Conveniently, we found that within this range, conditions can be selected based on the requirements of the DNA nanostructures or other system components.

DNA origami buffers are broadly compatible with liposome binding. DNA origami nanostructures often require specific ionic buffer conditions. Divalent cations such as Mg^2+^ stabilise DNA duplexes during nanostructure folding (72) and increase stability by inhibiting the electrostatic repulsion between DNA strands (54, 67). Some DNA nanostructures are designed to change shape in response to changes in ion concentration and pH, acting as sensors (18). While changes in external buffer can cause changes in liposome membrane density and diffusivity (56) (57), within the ranges tested here, addition of Na^+^ (0 – 200 mM) and Mg^2+^ (0 - 40 mM), resulted in only a small decrease in DNA-lipid bindings, and no increase in non-specific binding of unmodified DNA was observed for any ionic condition.

Both DOPE/DOPC and DPhPC lipid mixtures were found to work well for lipid binding of DNA strands. DNA is highly negatively charged, while these liposomes are zwitterionic. Liposomes with DOPE would be expected to be positively charged below pH 3.5 and negatively charged above pH 8, while those with DPhPC would be expected to be neutral in this pH range and only become charged at a very low pH (e.g. pH 2) (73). We observed no change in lipid-binding yield or non-specific binding across the range pH 4-10 for both DPhPC or DOPE/DOPC liposomes, which suggests lipid ionisation does not play a significant role in these conditions. However, we found that cholesterol-mediated lipid binding of DNA strands was completely inhibited in acidic conditions (pH = 2). Hydronium ions (H_3_O^+^) are known to promote lipid-lipid binding interactions within a bilayer (56) and affect the behaviour of water molecules at the membrane-water interface (57). This could potentially lead to the inhibited binding which was observed in strongly acidic conditions (72).

Interestingly, we found that increasing the cholesterol content of lipid mixtures above 20% increased the binding of DNA strands to DOPE/DOPC liposomes but slightly decreased binding to DPhPC liposomes. Cholesterol induces dense packing of phospholipids, reducing liposome permeability and increasing stability (74). Cholesterol also stabilises the structure of some membrane proteins and promotes highly curved membrane intermediates during fusion (75). Branched chain lipids like DPhPC occupy a greater area per molecule within a bilayer compared to linear-chain lipids such as DOPE and DOPC (76), and DPhPC has a lower cholesterol saturation limit than DOPE (77). Thus, the increased lipid binding of DNA strands that we observed with increasing cholesterol on DOPE/DOPC liposomes could be due to increased stability of the DNA-conjugated cholesterol in the membrane. The lower cholesterol saturation limit of DPhPC could explain why no further increase in lipid-DNA binding was observed above 20% cholesterol on DPhPC liposomes.

For small DNA motifs, dsDNA bound more efficiently to liposomes than ssDNA, and the addition of a 6-nt ssDNA overhang had minimal effect. Membrane-bound ssDNA has been observed to lie close to the surface of lipid bilayer membranes in fluorescence studies (78), while dsDNA remains in a stable position protruding normal to the membrane surface (79). Thus, the orientation difference of dsDNA compared to ssDNA may result in improved binding. The addition of a 6-nt overhang on cholesterol-tagged dsDNA strands has been proposed to assist nanostructure assembly by inhibiting strand aggregation (41). Inclusion of a 6-nt overhang next to the cholesterol group resulted in no significant decrease in binding for DOPE/DOPC liposomes, and but a significant, but small (△C_R_ = 0.10 ± 0.46) decrease in binding with the added overhang for DPhPC liposomes. This suggests that there is no large penalty from routine incorporation of overhangs on membrane-targeting nanostructures, but possible effects arising from other lipid compositions may need to be considered.

Upon studying the membrane binding of DNA strands, we next evaluated the membrane binding behaviour of our DNA origami tile. We measured the number of cholesterols present on our DNA origami tile using fluorescence labelling and observed a non-linear increase in fluorescence as the number of cholesterols was increased. This non-linear increase in fluorescence could possibly be due to non-linear range of detection provided by the gel analysis (80) or self-quenching of the Cy5 fluorophores in close proximity on the tile (81, 82), or from incomplete occupation of the H1 handles by cholesterol-strands (83). Assuming incomplete occupation and linear fluorescence intensity per handle for low numbers (n = 0-4), we are able to estimate a lower bound for the actual cholesterol number in the n = 16 sample. Linear regression across 0 to 4 handles estimates that our occupancy is 69% (for 16 cholesterols, our occupancy was 11 cholesterols with [7,21] at 95% CI). This closely matches previous reports for mean staple occupancy of DNA origami tiles, which was measured as 72% for the same 10x staple excess folding conditions (83).

The number of cholesterol groups on a DNA origami tile had a large effect on binding yield. Adding a large number of hydrophobic groups has been shown to be necessary for overcoming the energy penalty associated with pore formation (9), and beneficial for maintaining stable insertion of a transmembrane nanopore (15). However, a large number of cholesterol groups may inhibit the membrane-binding function of cholesterol-tagged DNA nanostructures by inducing aggregation (41) or structural deformities (37). We found in our gel electrophoresis measurements that a global optimum for membrane binding occurred with there were n = 4 cholesterols. However, this was dependent on which assay was used: an optimum number of cholesterols of 8 was observed with the gel-shift assay and an optimum number of 4 was observed with the microscopy.

We suggest that our observation of an optimal cholesterol number in these experiments is due to intra-tile and inter-tile transient binding of cholesterols. List and colleagues (37) showed that hydrophobic interactions between a large number of cholesterols (i.e. 35 cholesterols) on the DNA tile can result in the folding of the tile. Here we did not observe deformation or aggregation of the tile on folding with higher cholesterol numbers (Supplementary Figure 12, 13). However, it is possible that transient hydrophobic interactions occur between the cholesterol groups, which are not detected by gel or TEM. This would have the effect of decreasing the availability of cholesterols for binding to liposomes, and explain the observed decrease in membrane binding with increasing cholesterol number.

The difference in optimal cholesterol number obtained from the gel-shift assay and the microscopy is likely due to the different tile purification methods used. For the gel-shift assay, the tiles were gel purified. For microscopy, the tiles were purified by PEG precipitation to achieve higher concentrations of tile. PEG precipitation results in more aggregation compared to gel purification (Supplementary Figure 22). Higher aggregation results in more interaction between the tiles, and in turn, is expected to decrease membrane binding.

We found that positioning cholesterols along the edge of tile resulted in more binding compared to cholesterols positioned at the centre of the tile, consistent with a previous study (30). We also found that the membrane binding of tiles increases as the spacing between the cholesterols and the tile increases. These trends are likely due to greater accessibility of the cholesterols at larger distances from the tile, and located at the edges of the tiles. The tile is flexible (40) and can fold, and the centre of the tile is more likely to be hidden compared to the edges. This may limit the accessibility of the cholesterols positioned at the centre and result in decreased membrane binding.

For all the results above, some non-specific binding was observed with the control tile (no cholesterol) in the gel-shift assay. However, the amount of non-specific binding varied from gel to gel, and was found to vary between preparations of SUVs. Percentages of no-cholesterol tiles bound to the SUVs ranged from 15% to 60% across all the gels run in all different experiments. To control for this, our samples were always compared within gels, not between gels. Generally, microscopy experiments had smaller error ranges, while gel assays had larger errors but were higher throughput and useful for comparing large numbers of conditions.

We found that toehold-mediated strand displacement could be used to remove cholesterol modified strands from the DNA tile with high efficiency, but that displacement efficiency decreased if the cholesterol was positioned directly adjacent to the toehold. This is likely due to the interaction of the toehold with the cholesterol group when it is in proximity. Previous work by Ohmann and colleagues showed that an overhang placed next to a cholesterol can interact with it to reduce aggregation (41). An increase in temperature from room temperature to 37°C increased the displacement efficiency of these proximal designs, to a similar value as designs where the cholesterol was located distal to the toehold.

Toehold-mediated strand displacement was shown to remove DNA tiles from liposomes, by separating the tile from the cholesterol strand. In this case, the cholesterol-modified DNA strand is expected to remain docked to the liposome. For cholesterol displacement of tiles already bound to liposomes, displacement was efficient for cholesterols positioned both proximal and distal to the toehold. This is in contrast to strand displacement in the absence of liposomes, where proximal toeholds had reduced efficiency. When the tile is bound to the liposome, we expect the cholesterol to have inserted into the bilayer. This may result in less interaction between cholesterol and toehold, making a more accessible toehold, facilitating more efficient strand displacement (Supplementary Figure 23). We expected to see reduced displacement efficacy for designs with shorter linkers (e.g. Dt1.4 compared to Dt15.2, Pt8.5), because tight binding of the tile to the liposome might sterically hinder strand displacement. However, results here were dominated by the strong effect of linker length on initial binding yield. The short linker design had very low initial yield, comparable to non-specific binding, and so it was not possible to detect if there was a decrease on strand displacement with the gel shift technique.

## CONCLUSION

In this work, we tested different lipid species and DNA configurations to screen for optimal conditions to promote DNA-lipid binding. Our results suggest that lipid type, pH and DNA configuration are the most important parameters to consider when optimising for the binding of DNA strands to liposomes, whereas mono- and divalent-salt concentration play a minor role.

Our results have shown that the membrane binding of DNA nanostructures to liposomes can be optimised by changing the cholesterol number, cholesterol configuration and cholesterol distance from the DNA nanostructure. We found that the optimal number of cholesterols for membrane binding of a 2D DNA origami tile is between 4 and 8, and that membrane binding is more favourable when cholesterol groups are placed at the edge of the tile compared to the centre of the tile. A larger linker length between the tiles and the cholesterol also results in greater membrane binding.

We demonstrated reversible membrane binding of the DNA nanostructures onto liposomes using toehold mediated strand displacement. The efficiency of strand displacement is reduced if the toehold is adjacent to the cholesterol in unbound DNA nanostructures, but not for lipid-bound DNA nanostructures.

Future work could extend the findings from this work to more complex 3-dimensional DNA nanostructures with greater functionality. There is a general trade-off between increasing the lipid binding yield and decreasing aggregation. The flexibility of the 2D tile plays a role in aggregation, and so the greater rigidity of 3D DNA origami nanostructures may be an advantage. We anticipate our findings will provide guidelines for the design of more complex membrane binding DNA nanostructures with broad applications in nanomedicine, nanotechnology, and nucleic acid research.

## Supporting information

Supplementary Table of Nucleotides

