## Supplementary Methods, Supplementary Table 1, Supplementary Figures 1-23 for "Binding of DNA origami to lipids: maximising yield and switching via strand-displacement"

### Supplementary information

#### Supplementary Methods

##### Lipids and oligonucleotides

DNA strands were visualised by the addition of a 5' Alexa 647 dye. Hydrophobic modifications of DNA strands were made by adding a 3' cholesterol moiety attached via a tetraethylene glycol spacer (TEG-cholesterol) (Integrated DNA Technologies, Inc., USA). DNA was suspended at 100 µM in MilliQ water. Fluorescent DNA was stored at –20°C in 20 µL aliquots covered in foil.

Sequences for DNA strands, duplexes and fluorophores were generated using the NUPACK design software (3) and tested to avoid the formation of unwanted secondary structures. The sequence for the 6 nt overhang added to the 5′ end of a strand complementary to the fluorescent cholesterol strand, was taken from a previously characterised design (4).

Liposomes were made from two main lipid mixtures: a 1:1 binary mixture (weight:weight) of DOPE and DOPC and 100% DPhPC. Tethering of liposomes to a cover slip surface was enabled by adding 0.1% (weight) PE-biotin. Fluorescent imaging was enabled by adding 0.1% (weight) PE-Rhodamine or by encapsulating red fluorescent protein (mRFP1). Lipids (all from Avanti Polar Lipids Inc., USA) were suspended in chloroform at 10 mg/mL and stored at -20°C. Unless otherwise stated, all descriptions of lipid concentration are stated in terms of mass.

##### Production of small unilamellar liposomes by extrusion

Lipid stocks were added to a round-bottom glass tube and dried into under nitrogen into a thin film and resuspended in extrusion buffer (210 mM sorbitol, 100 mM NaCl, 5 mM Tris-HCl, pH 7.5) to final concentration of 1 mg/mL by vortex mixing and sonication. The suspension was transferred to a 500 µL glass syringe (Hamilton Company, UK) and passed back and forth through a 100 nm polycarbonate filter (Whatman plc, USA) using a Mini-Extruder kit (Avanti Polar Lipids Inc., USA) 41 times to produce a clear suspension of liposomes.

##### Production of giant unilamellar liposomes by electroformation

Giant unilamellar liposomes (GUVs) were prepared by electroformation using the Vesicle Prep Pro machine (Nanion Technologies GmbH, Germany). 30 µl of 3.5 mg/mL lipid dissolved in chloroform was added to a conductive indium tin oxide-coated glass slide and spread over a spot approximately 12 mm in diameter and allowed to air-dry for two minutes into a circular film. A 1.5 mm thick rubber gasket of 15 mm diameter was placed around the film, forming a well with 250 µl of electroformation solution (210 mM sorbitol, pH 7.5). A second indium tin oxide-coated glass slide was placed face-down on top of the gasket and clamped in place, creating a sealed chamber of liquid between the two slides. The machine was run using the default protocol of 3 V AC for 120 minutes.

**The effect of detergents on DNA-liposome colocalisation**

The tolerance of DOPE/DOPC liposomes and DPhPC liposomes to polysorbate 20 detergent was tested by incubating liposomes in 210 mM sorbitol, 100 mM NaCl, 5 mM Tris-HCl, pH 7.5 buffer with various concentrations of polysorbate 20 for one hour (Supplementary Fig. 1).

Liposomes were produced in buffer solution consisting of 210 mM sorbitol, 100 mM NaCl, 5 mM Tris-HCl, pH 7.5. Buffer solutions of the same composition plus up to 0.5% (volume) polysorbate 20 were then prepared and used to flush excess streptavidin from slides prior to the addition of liposomes. The same buffer was also used to dilute liposomes and DNA prior to their addition to the slide, to ensure the concentration of detergent remained stable.

Low colocalisation ratios were measured for all DNA configurations in buffers containing 0.1% detergent and above, or all values above the CMC of polysorbate 20 detergent (0.066 %vol) (1).

**Threshold determination for co-localisation analysis**

For image colocalisation analysis, images of liposomes were converted into a binary image. To accomplish this, pixels of intensity values greater than a certain threshold were determined to be 'liposomes' and pixels of intensity values below this threshold were determined to be 'background'. This threshold was then applied to divide the image into two sections: 'liposomes' and 'background'. In order to determine a suitable pixel intensity threshold for dividing an image of liposomes into two sections, we tested the following formula:

$$T = \bar{x} + ks$$

Where $T$ is the threshold of pixel intensity applied to divide the image into two sections, $\bar{x}$ and $s$ are, respectively, the mean and standard deviation of pixel intensity, and $k$ is the multiplier of standard deviation (ranging from 1-4, Supplementary Fig. 2). Analysing the same image with k set from between 1 to 4 determined that k = 2 was best suited to mask for liposomes by intensity. With k = 2, the selection included all visually distinguishable liposomes, but did not create larger, diffuse, non-circular selections around larger liposomes.

**Colocalisation ratio variation with liposome coverage**

Conventional colocalisation metrics for biological image analysis such as Pearson's Correlation Co-efficient and Manders Overlap are affected by the amount of background included in the analysed area, that is, reported colocalisation values vary with the proportion of unlabelled dark regions. Accordingly, these metrics are typically used to describe the distribution of two analytes within a defined region such as a cell after the image background has been removed (2). We developed a custom analysis procedure to quantify the colocalisation of DNA and liposomes in two-colour images (Supplementary Fig. 3) and observed that in changing conditions the liposome coverage of the surface could differ (Supplementary Fig. 4). We confirmed that our metric was not affected by the relative area of each section by plotting liposome area against colocalisation ratio and observing no trend in colocalisation ratio that was dependent on liposome coverage (Supplementary Fig. 5). Measuring as a ratio, rather than the mean pixel intensity of DNA on liposomes, accounts for minor variations in the overall amount of DNA added to each slide.

**Fluorescence measurements of ssDNA and dsDNA in different buffer conditions**

DNA duplexes were annealed at 10 µM final concentration in duplex buffer (100 mM NaCl, 5 mM Tris-HCl, pH 7.5). Oligos were heated to 90˚C for five minutes then cooled in a thermocycler at 5˚C /minute for 15 minutes to 15˚C, and stored at 4˚C. For dsDNA assembly, unmodified complementary strands were added in a 3-fold excess to modified strands.

For fluorescence measurements, ssDNA and dsDNA were diluted to 100 nM in different buffer conditions as the following:

- pH experiments: 5 mM Tris-HCl, 210 mM D-Sorbitol, 100 mM NaCl (pH adjusted to 2, 4, 6, 7, 8, 10)
- NaCl experiments: 5 mM Tris-HCl, pH 7.5, 10 mM $MgCl_2$, 0 - 400 mM NaCl
- $MgCl_2$ experiments: 5 mM Tris-HCl, pH 7.5, 100 mM NaCl, 0 - 80 mM $MgCl_2$,

20 µL of sample was loaded into a 384-well microplate (6008260, Perkin Elmer) and the fluorescence intensity (635-20, 680-20) was measured using the PHERAstar FSX microplate reader (BMG Labtech). Three repeats of each measurement were performed (Supplementary Fig. 6).

### Supplementary Tables

| Abbreviation | Chemical Name | Mass |
|---|---|---|
| DOPE | 1,2-dioleoyl-sn-glycero-3-phosphoethanolamine | 717.531 g/mol |
| 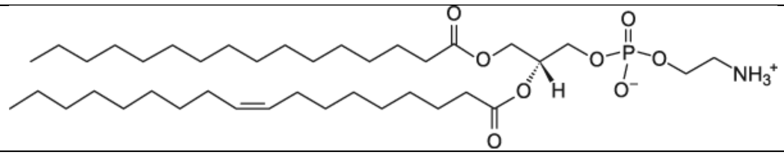 | | |
| PE-Rhodamine | 1,2-dioleoyl-sn-glycero-3-phosphoethanolamine-N-(lissamine rhodamine B sulfonyl) (ammonium salt) | 1300.712 g/mol |
| 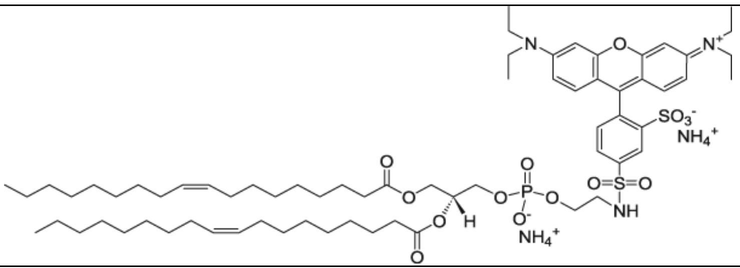 | | |
| PE-biotin | 1,2-dioleoyl-sn-glycero-3-phosphoethanolamine-N-(biotinyl) (sodium salt) | 991.606 g/mol |
| 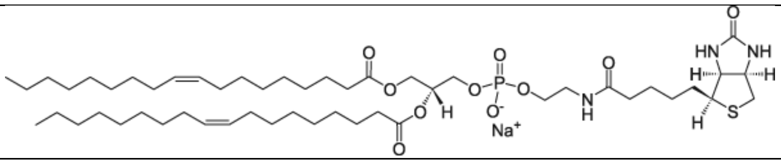 | | |
| DOPC | 1, 2-dioleoyl-sn-glycero-3-phosphocholin | 759.578 g/mol |
| 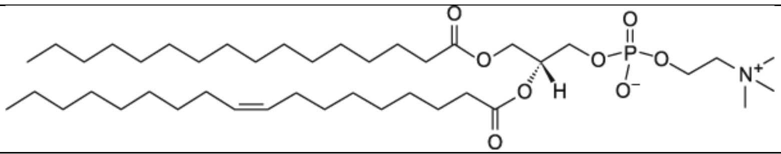 | | |
| DPhPC | 1,2-diphytanoyl-sn-glycero-3-phosphocholine | 845.687 g/mol |
| 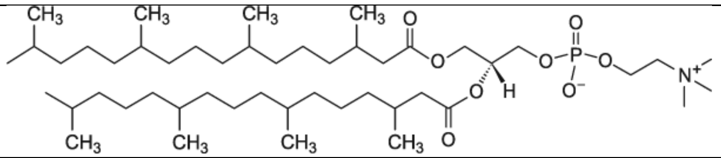 | | |
| Cholesterol | Cholesterol | 386.355 g/mol |
| 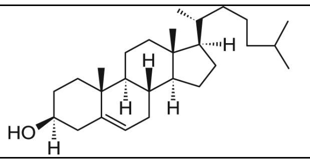 | | |

**Supplementary Table 1 – Lipid chemical structures**

Name, chemical structure and mass of lipid molecules used in liposome formation (5) .

| Reagent | Supplier |
|---|---|
| D-Sorbitol | S1876, Sigma |
| Tris-HCl | T3253, Sigma |
| NaCl | AJA465, Ajax-Finechem |
| $MgCl_2$ | AJA296, Ajax-Finechem |
| All DNA strands | Integrated DNA Technologies (IDT) |
| MilliQ water | Milli-Q, Millipore |
| DOPE 18:1 | 850725P, Avanti Polar Lipids |
| DOPC 18:1 | 850375P, Avanti Polar Lipids |
| DPhPC | 850356P, Avanti Polar Lipids |
| PE-rhodamine | 810150P, Avanti Polar Lipids |
| PE-biotin | 870282P, Avanti Polar Lipids |
| Cholesterol | 700000P, Avanti Polar Lipids |
| M13mp18 ssDNA | P-107, Bayou Biolabs |
| SyBrSafe stain | S33102, Thermo Fisher Scientific |
| PEG 8000 | P5413, Sigma |
| EDTA | E9884, Sigma |

**Supplementary Table 2 – List of reagents used**

**Supplementary Figures**

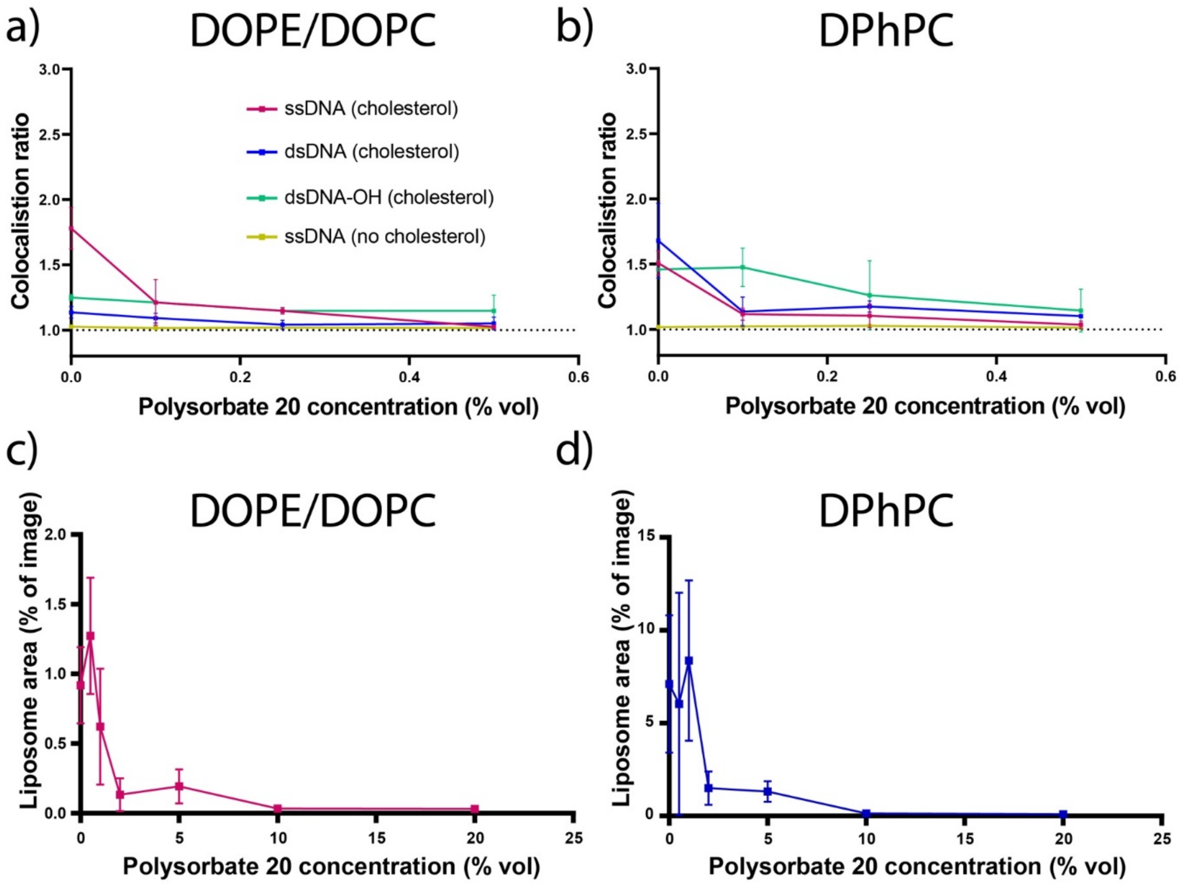


**Supplementary Figure 1 – DNA-Lipid colocalisation with detergent**

Liposomes and DNA colocalisation with increasing Polysorbate 20 concentrations. Colocalisation ratios (±SD) for DOPE/DOPC (A) and DPhPC (B) liposomes are shown for cholesterol-tagged ssDNA (magenta), cholesterol-tagged dsDNA (blue) and cholesterol-tagged dsDNA with a 6-nt overhang (green) and ssDNA with no cholesterol tag (yellow). Liposome survival is plotted at higher concentrations of Polysorbate-20, showing percentage of liposome area reduces to zero as detergent is introduced at high concentrations (>10%) for DOPE/DOPC (C) and DPhPC (D) liposomes.

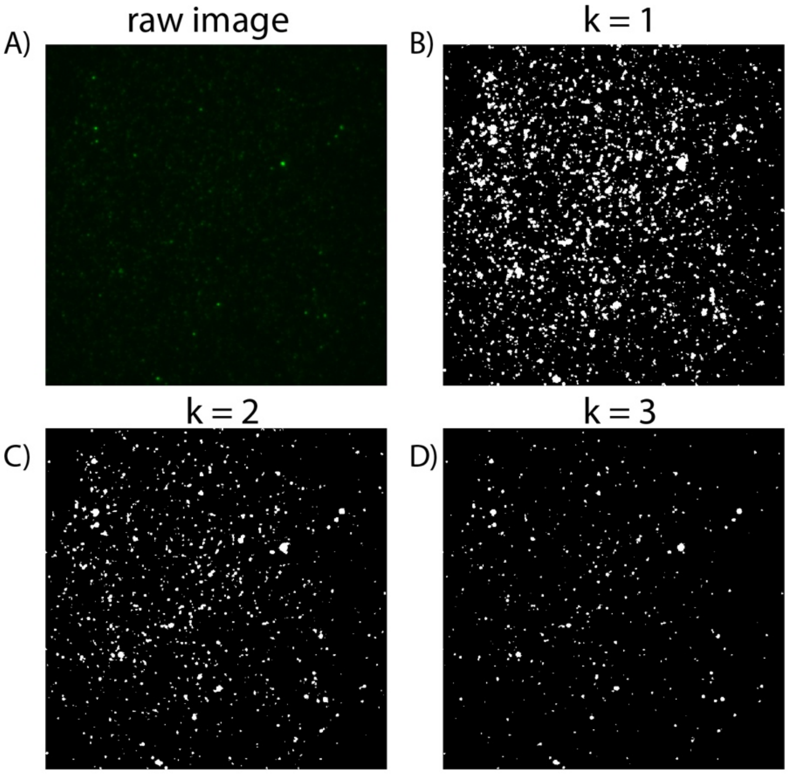


**Supplementary Figure 2 – Generating image mask**

Liposomes converted into a binary mask using three separate criteria A) Original image of liposomes (green) imaged in TIRF with 561 nm illumination. B) Mean plus one standard deviation. C) Mean plus two standard deviations. D) Mean plus three standard deviations. In each case, the image is divided into two section: ‘liposomes’ (white) and ‘background’ (black). The 2 standard deviation case was taken as the threshold that gave the best assignment of lipid and background pixels.

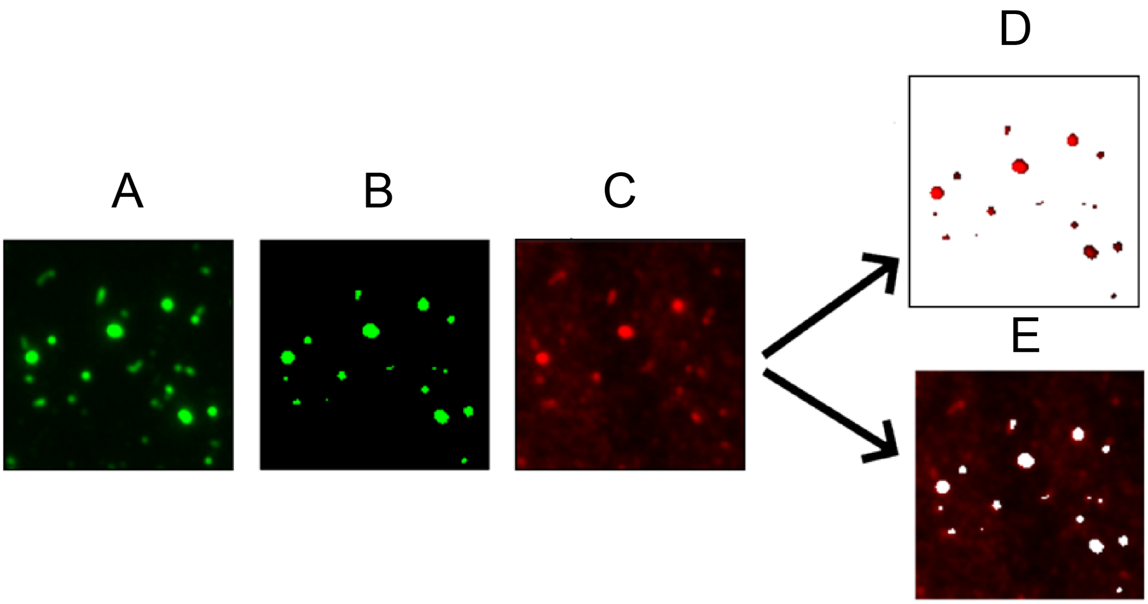


**Supplementary Figure 3 – Image analysis steps**

Colocalisation analysis procedure for quantifying DNA-liposome binding. (A) Liposome channel raw image. (B) Liposome channel converted into a binary image consisting of two sections, 'liposomes' and 'background'. (C) DNA channel prior to being split into two sections. (D) The 'liposomes' section of the DNA channel. (E) The 'background' section of the DNA channel. The mean pixel intensity in (D) is divided by that of (E) to calculate the colocalisation ratio. This number represents the relative intensity of DNA attached to liposomes compared to DNA found in the background. For example, a colocalisation ratio of $C_R$ = 1 indicates that DNA fluorescence is evenly distributed between liposomes and background, while $C_R$ = 2 indicates there is roughly twice as much DNA colocalised with liposomes compared to background

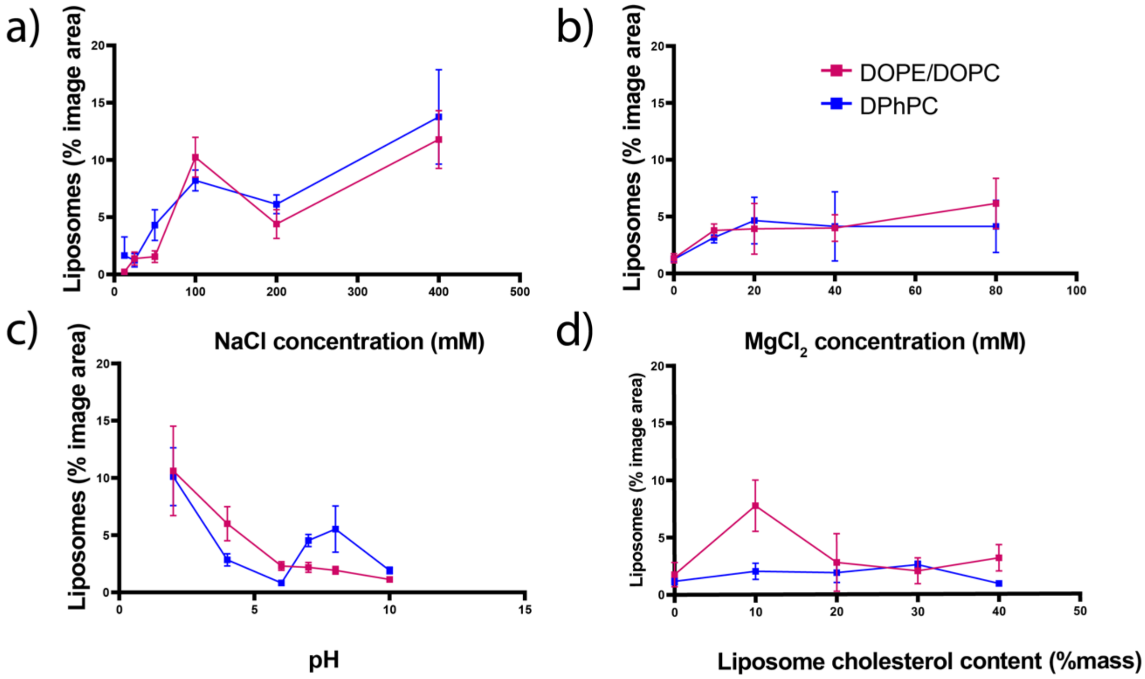


**Supplementary Figure 4 – Liposome image area**

Liposome variation with condition. Variation in liposome area (percentage of image area, mean ± SD) for varied (a) NaCl, (b) $MgCl_2$, (c) pH and (d) cholesterol content. Shown for DOPE/DOPC (red squares) and DPhPC (blue squares).

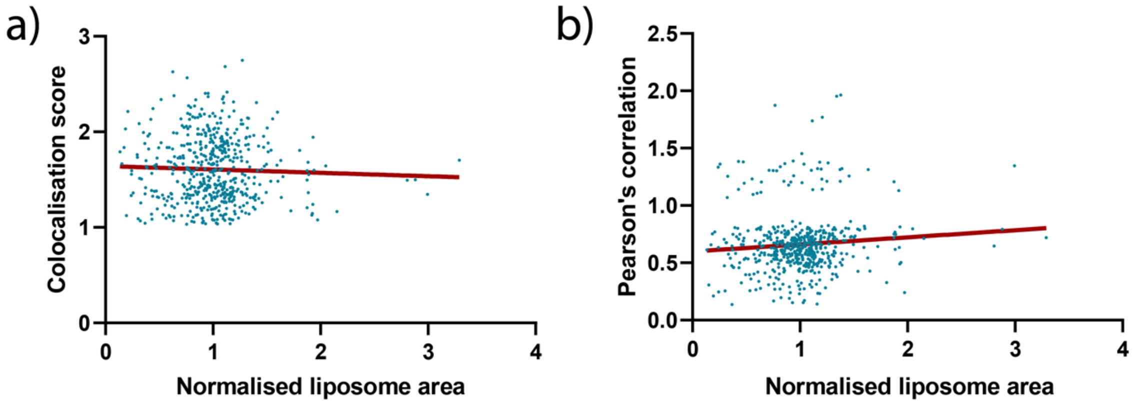


**Supplementary Figure 5 – Testing interdependence of colocalisation ratio and lipid area**

Scatter plot for all measured colocalisation ratios against normalised liposome area, with overlaid linear regression. All images containing cholesterol-tagged ssDNA (n=528) were analysed to test for correlation with liposome coverage in the image. Liposome area was normalised as follows: the area of the image occupied by liposomes (expressed as a percentage of pixels) for each individual image was divided by the mean of all twelve images for that buffer/DNA condition. Linear regression analysis of (A) colocalisation ratios against normalised liposome area and (B) Pearson's correlation coefficient against normalised liposome area was used to test for correlation. For colocalization score as used in this work, there was no correlation (95%CI gradient: 0.036 ± 0.079); for Pearson’s correlation coefficients against normalised area there was a non-zero gradient (95% confidence interval for slope = 0.062 ±0.060).

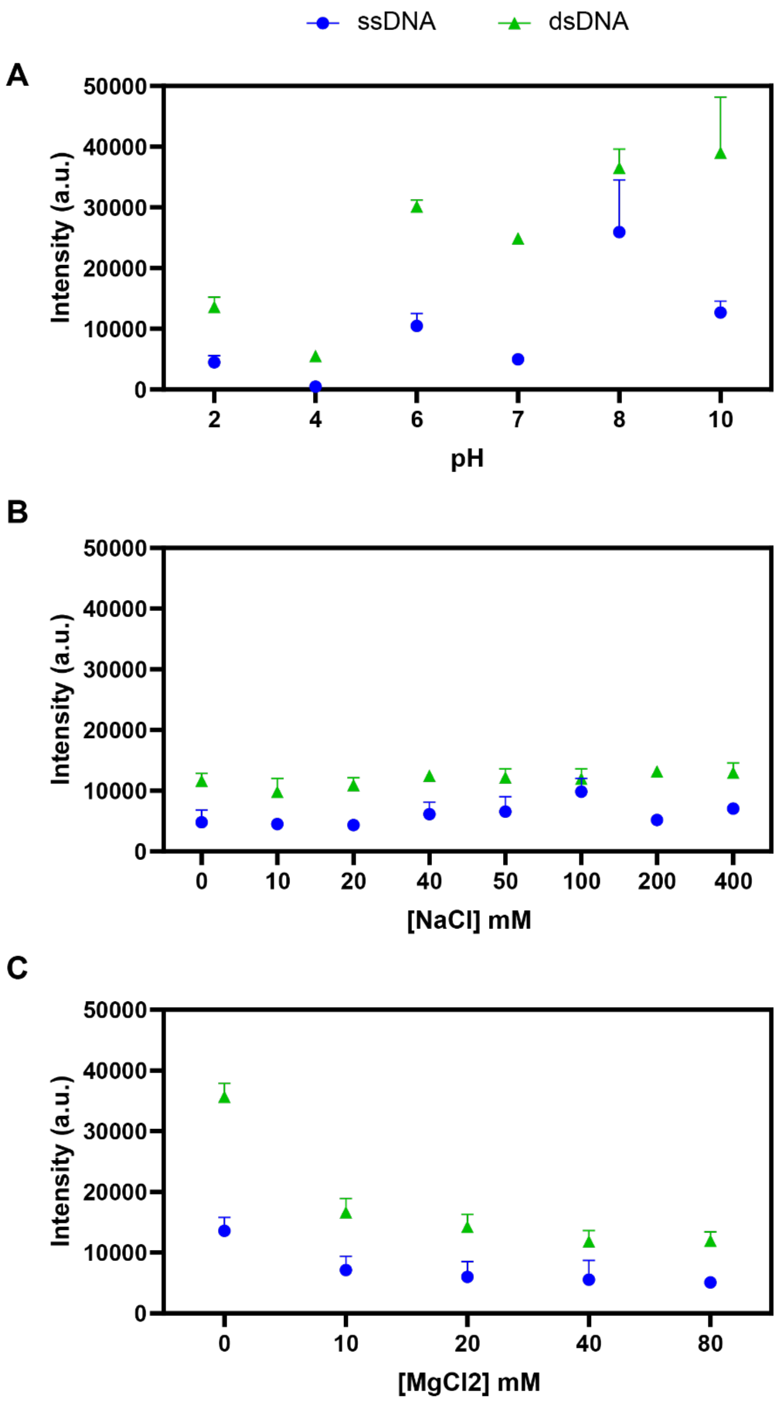


**Supplementary Figure 6 – Effect of buffer on fluorescence intensity**

The intensity of Alexa-647 labelled ssDNA and dsDNA in different buffer conditions.

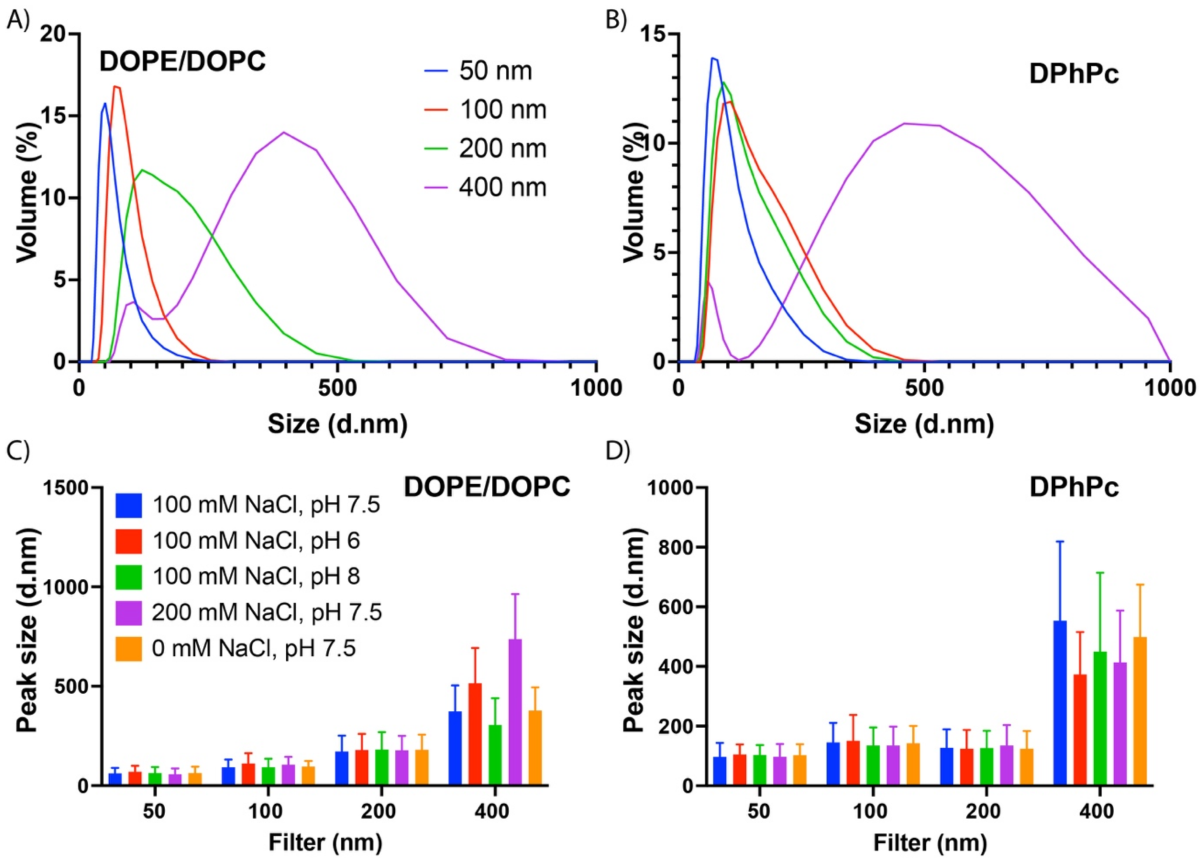


**Supplementary Figure 7 – Liposome characterisation by light scattering.**

Light scattering characterisation by zetaziser for 50 (blue), 100 (red), 200 (green) and 400 nm (purple) pore size extrusion filters respectively. Volume (%) is shown vs size for single measurement trace for each filter type for (A) DOPE/DOPC and (B) DPhPC liposomes with liposome buffer inside and outside the liposome (210 mM sorbitol, 100 mM NaCl, 5 mM Tris-HCl, pH 7.5). Data from mean of triplicate measurements indicating peak size (volume % vs size) with standard deviation reported as error bars for (C) DOPE/DOPC and (D) DPhPC liposomes. For separate characterisation here, internal composition of liposomes was fixed across all measurements (liposome buffer), while external conditions were varied: all with 210 mM sorbitol, 5 mM Tris-HCl; 100 mM NaCl pH 7.5 (blue, liposome buffer); 100 mM NaCl pH 6 (red); 100 mM NaCl pH 8 (green); 200 mM NaCl pH 7.5 (purple); 0 mM NaCl  pH 7.5 (yellow).

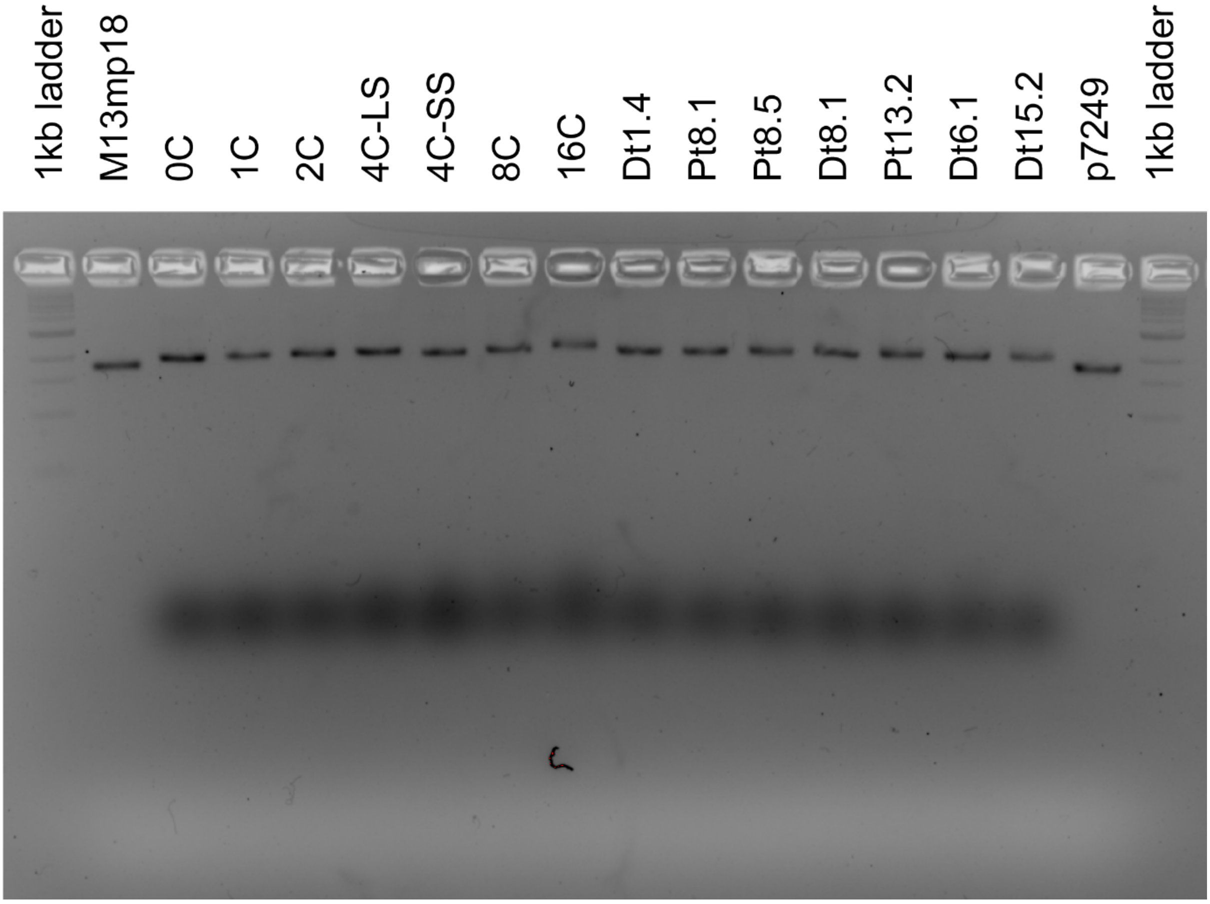


**Supplementary Figure 8 – Agarose gel analysis of DNA tiles**

Gel image for all the DNA tiles used in this work. For simplicity, all tiles here were folded without any fluorophores. 1kb ladder and the p7249 scaffold are loaded as markers. The gel is a 2% agarose gel run for 2.5 hours at 60 V at RT. All subsequent gels presented in this SI were also run at similar conditions.

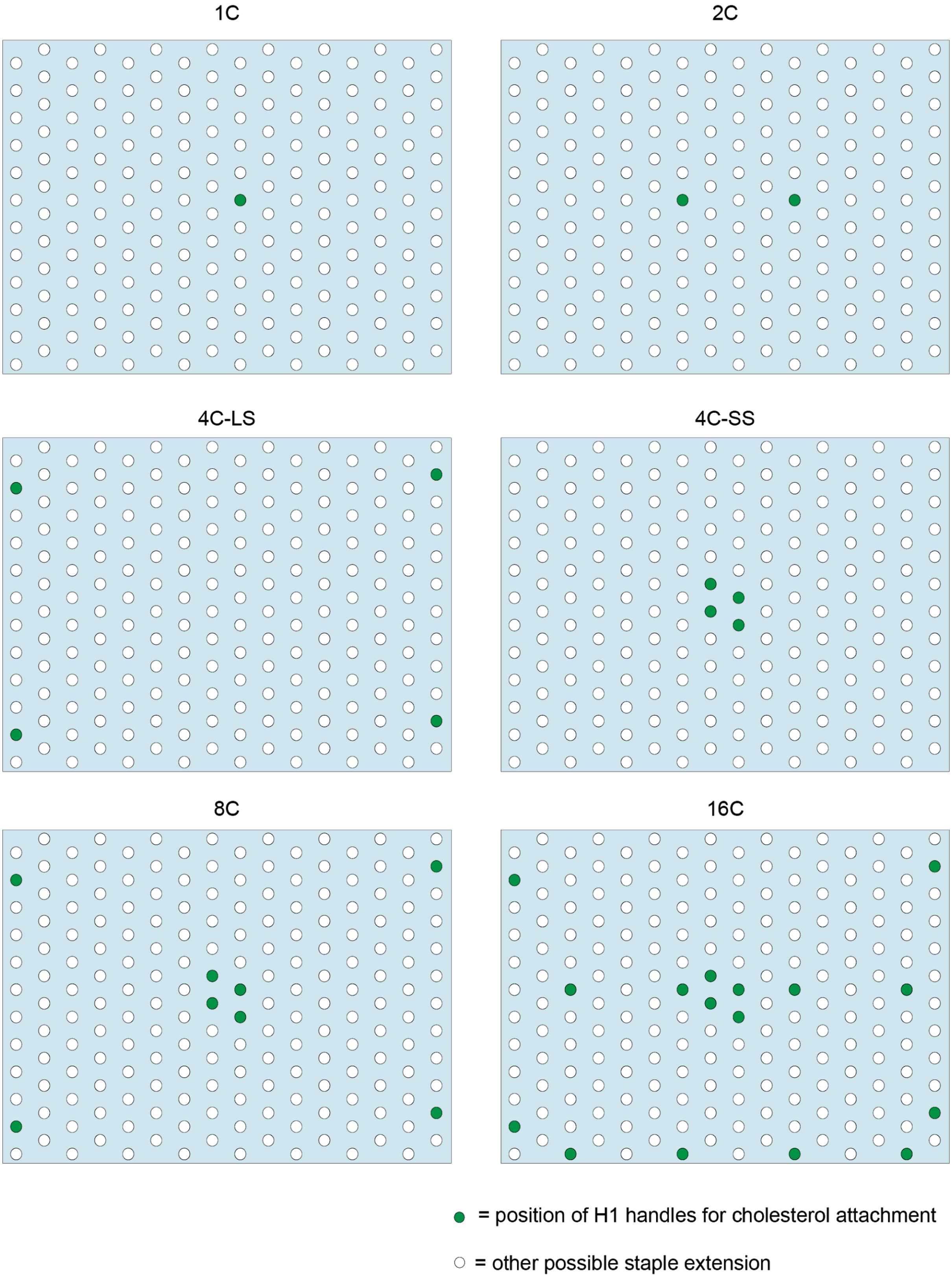


**Supplementary Figure 9 – DNA tile H1 handle layouts**

Schematic showing the positions of H1 handles (green circles) for cholesterol attachment. White circles represent possible staple extensions across the tile. A simpler schematic of this Figure is given in Figure 4A.

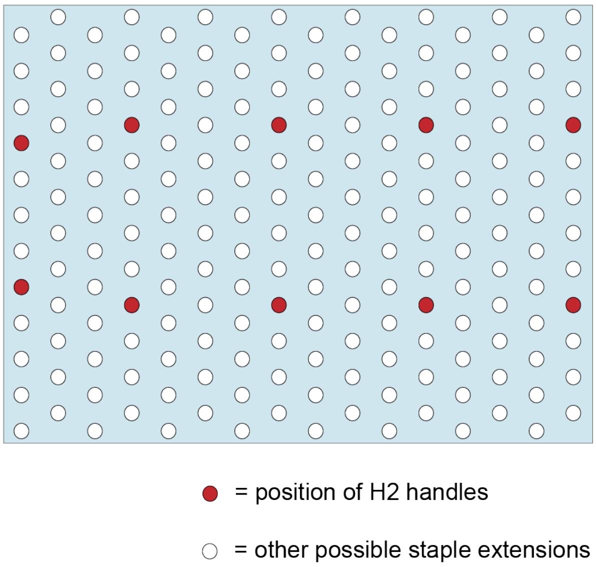


**Supplementary Figure 10 – DNA Tile H2 handle layout**

Schematic showing the positions of H2 handles (red circles). White circles represent possible staple extensions available across the tile.

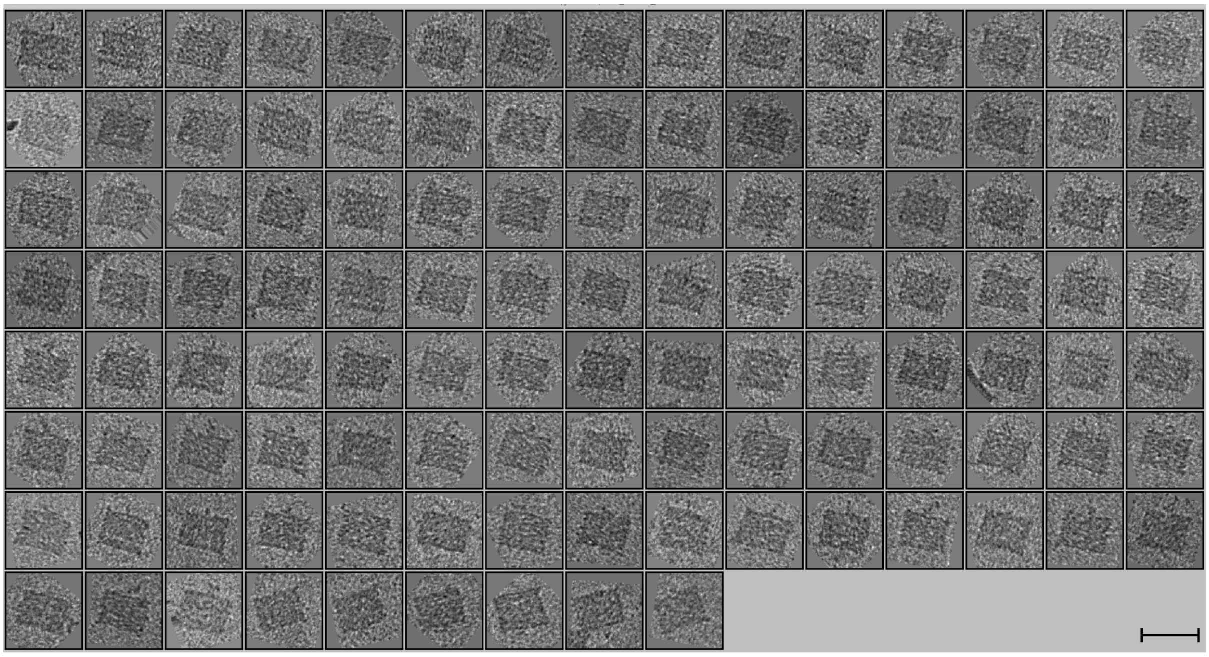

**Supplementary Figure 11 – TEM analysis of DNA tile**

114 representative particles of the DNA tile. These particles were picked manually and averaged using RELION 3.0.6 (6) to give the averaged particle in Figure 1.A. Scale bar: 100 nm.

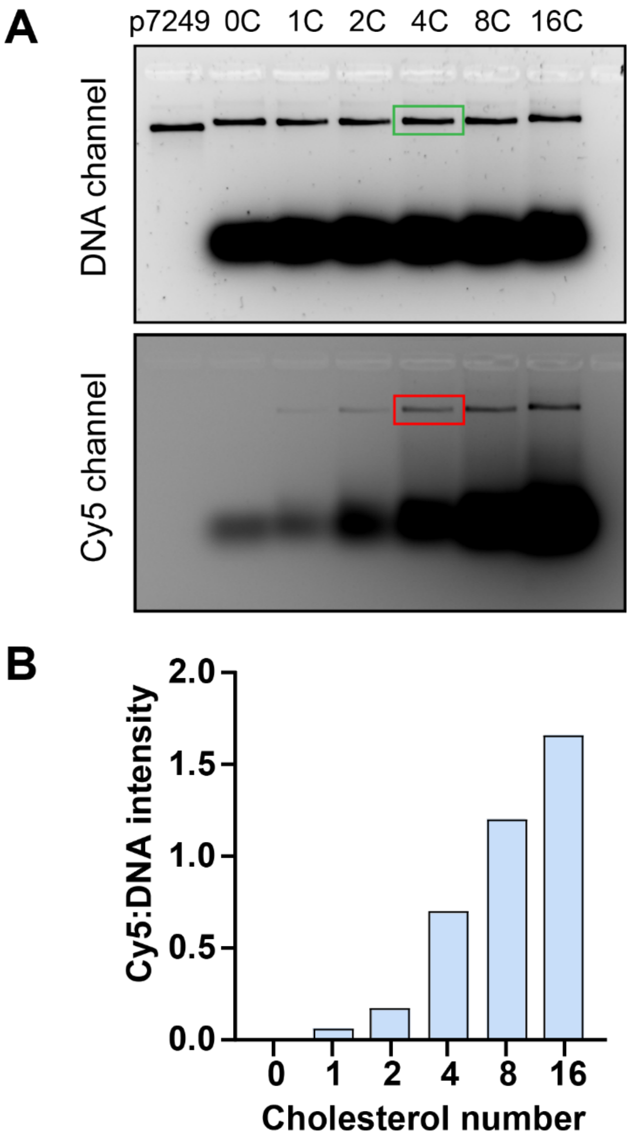


**Supplementary Figure 12 – Validation of cholesterol attachment to the tile**

Validation of cholesterol attachment to the tile for different cholesterol numbers. (A) Gel image. DNA and Cy5 channels which represent the tile and Cy5 fluorophore, respectively are shown. The integrated band intensity in the Cy5 channel (red box) is divided by the integrated band intensity in the DNA channel (green box) to obtain the ratio of Cy5:DNA intensity. This is tabulated in the chart in (B).

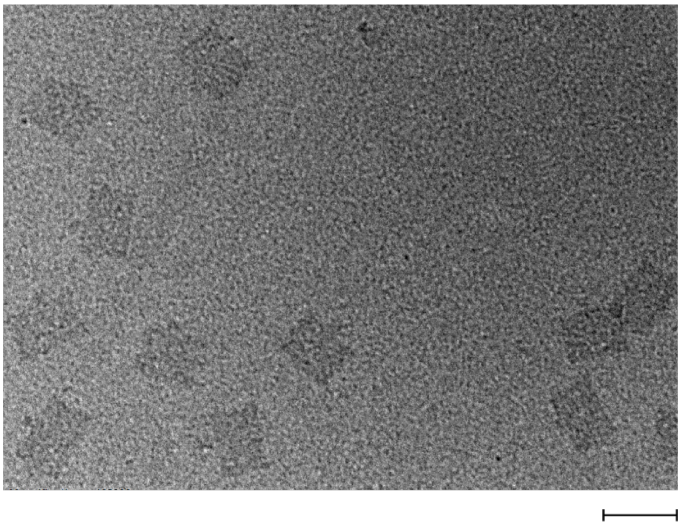

**Supplementary Figure 13 – TEM image of DNA tile with 4 cholesterols**

TEM image of the DNA tile folded with 4 cholesterols. Scale bar: 100 nm

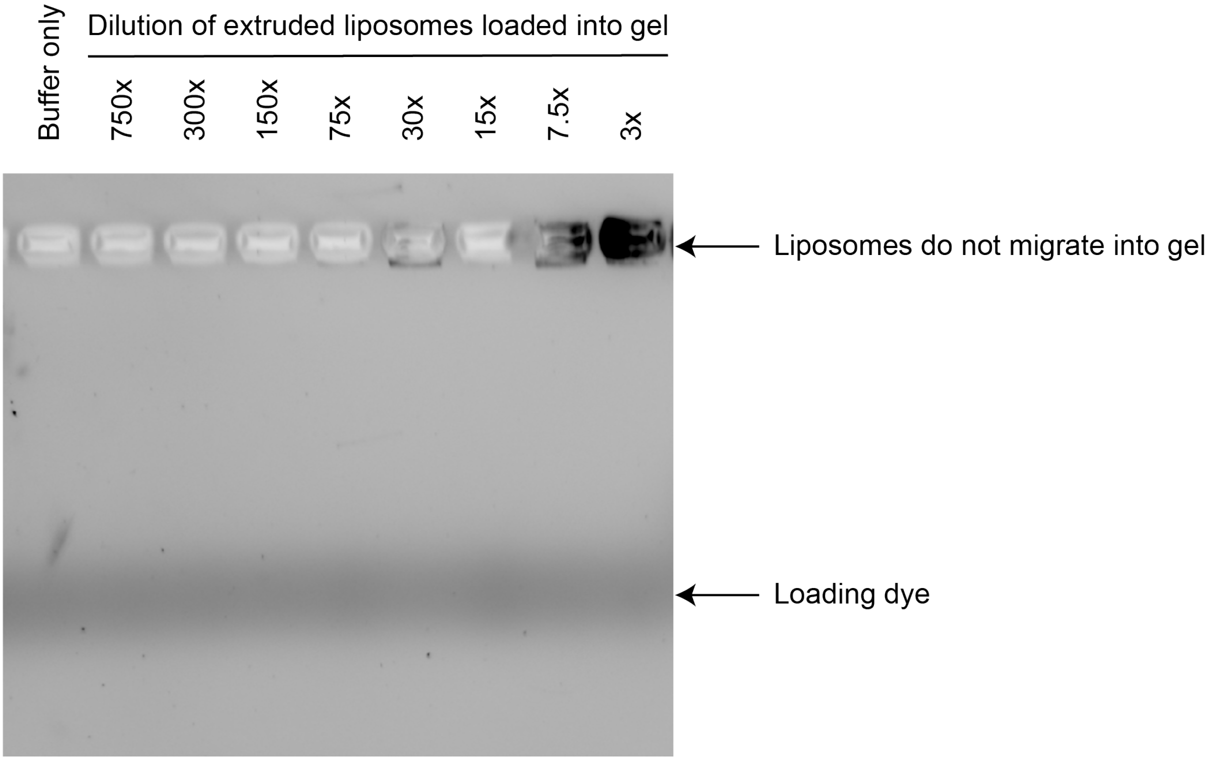


**Supplementary Figure 14 – Agarose gel analysis of extruded liposomes**

Gel image showing that rhodamine labelled extruded liposomes do not migrate into the gel. Gel was imaged using the Epi-green, 520–545 nm excitation and 577–613 nm filter. The liposomes were diluted X-fold (annotated above each well) prior to loading into gel. As the dilution of the extruded liposomes decreases (higher concentration), the band in the well becomes more prominent. No extruded liposomes were observed in the gel. A buffer only control is also shown in lane 1.

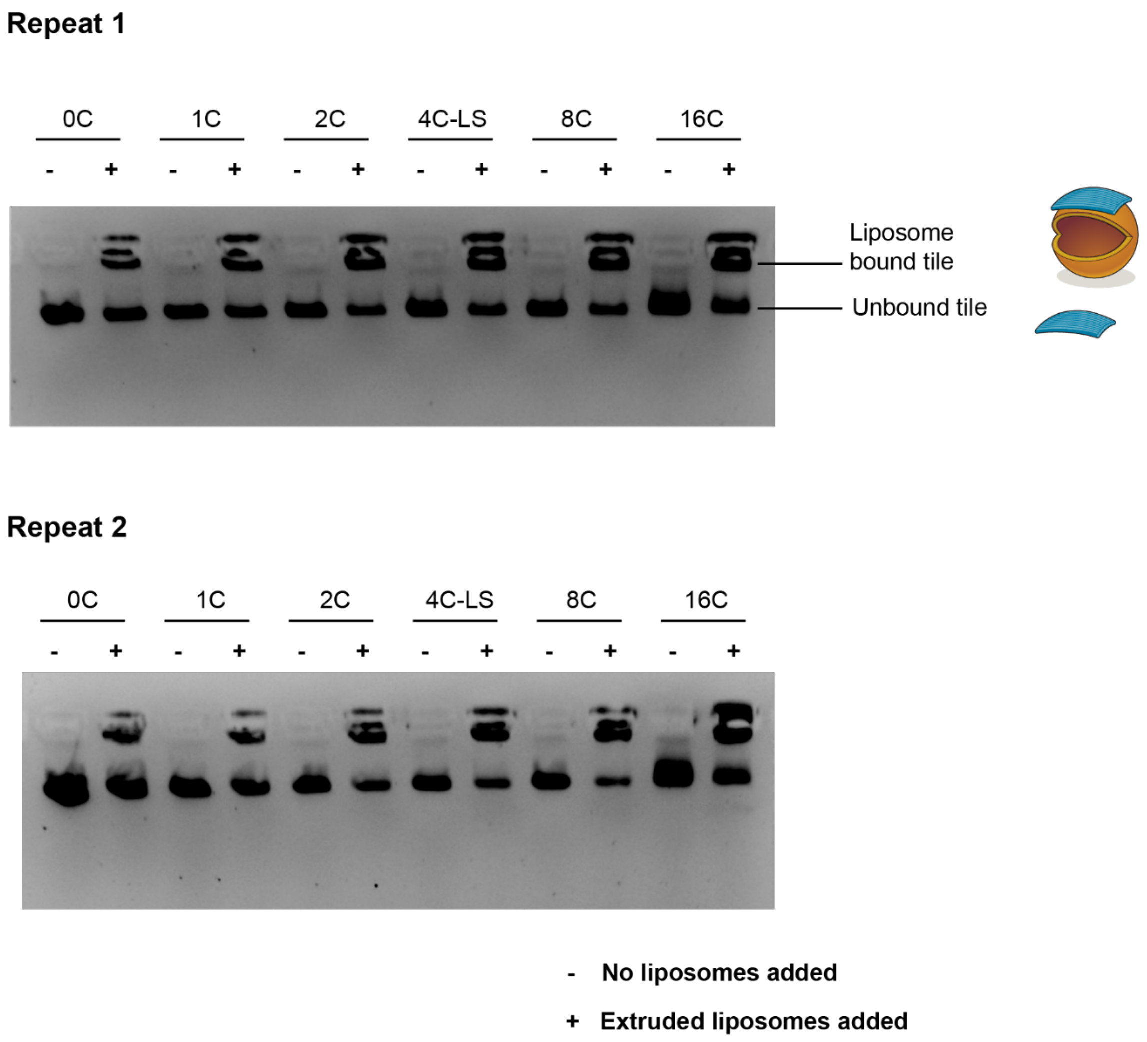


**Supplementary Figure 15 – Membrane binding versus cholesterol number, repeat gels**

Gel-shift assay for membrane binding with different number of cholesterols on the DOT. Two repeats were performed.

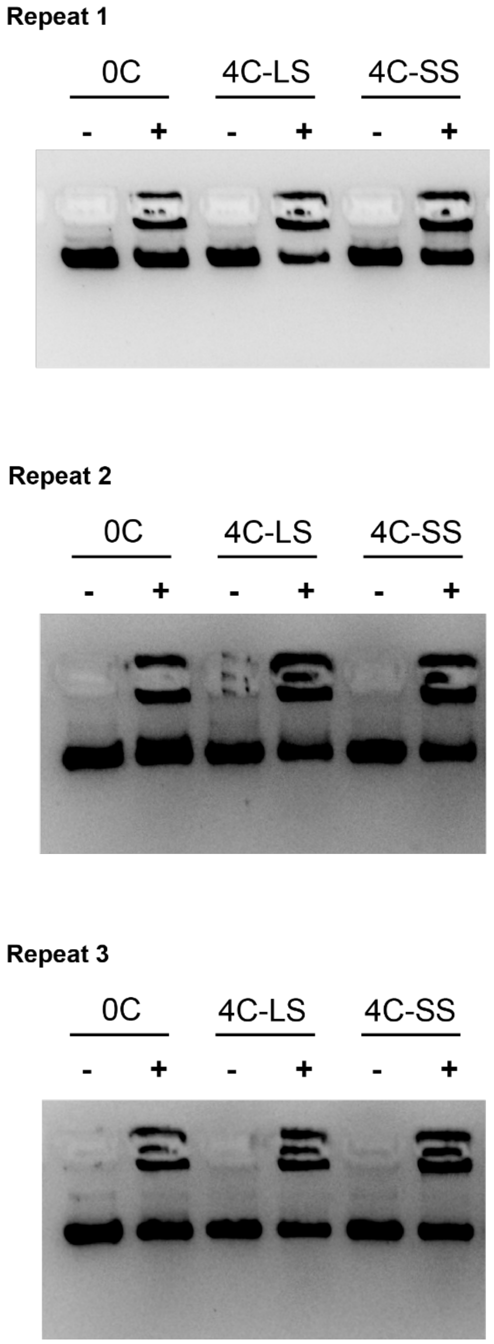


**Supplementary Figure 16 – Membrane binding versus cholesterol configuration, repeat gels**

Gel-shift assay for membrane binding with different configuration of cholesterols on the DOT. Three repeats were performed.

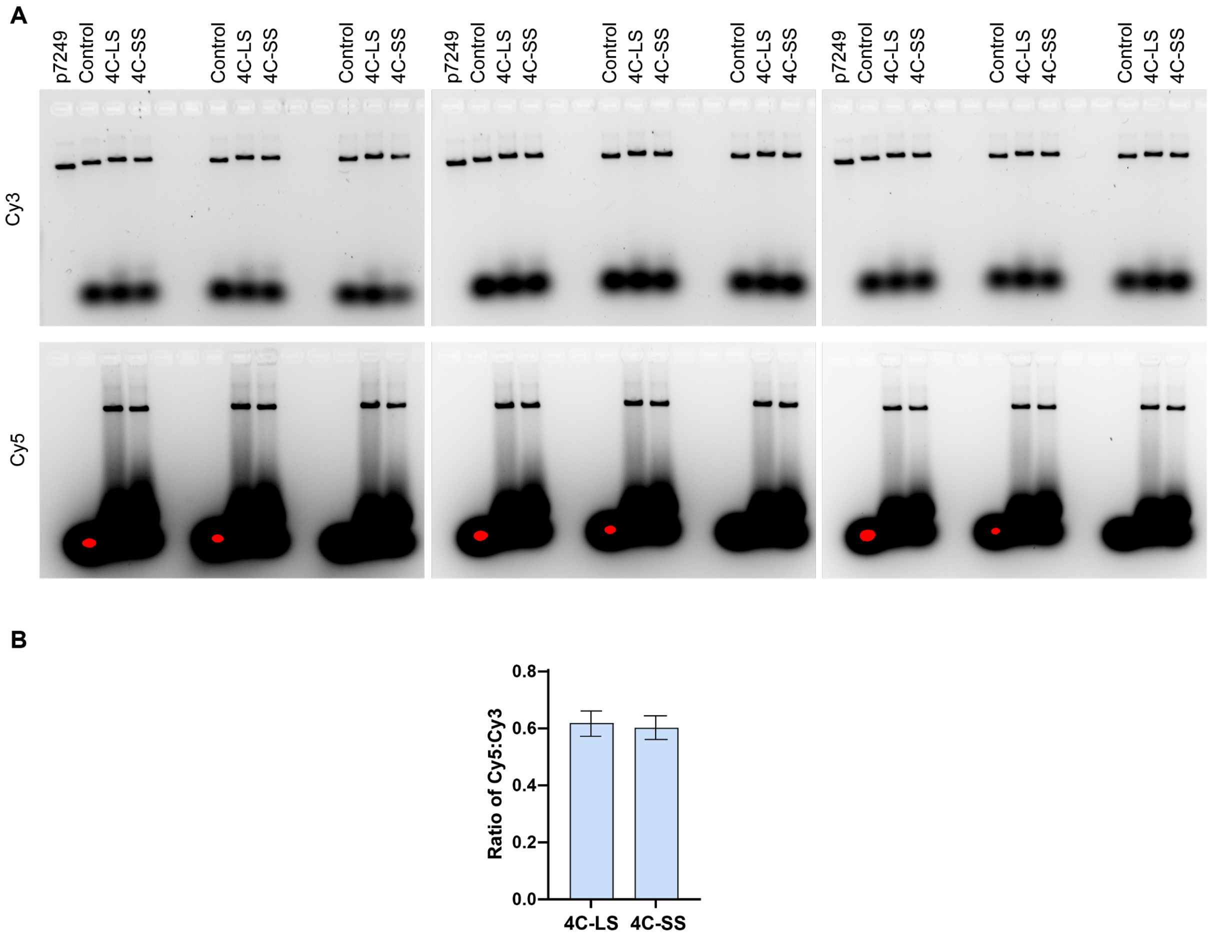


**Supplementary Figure 17 – Number of cholesterols per tile, repeat gels**

A. Gels showing the attachment of cholesterols (Cy5 channel) to the tile (Cy3 channel) for the large square (4C-LS) and small square (4C-SS) configurations. Nine repeats were done across three gels. The control is the tile with no cholesterols. B. Cy5:Cy3 ratio as tabulated from the gels in A. The differences were not significant. The cholesterol attachment is similar for both configurations. Red spots indicate camera saturation for unincorporated excess DNA origami staples from the folding reaction, which are not relevant to this analysis.

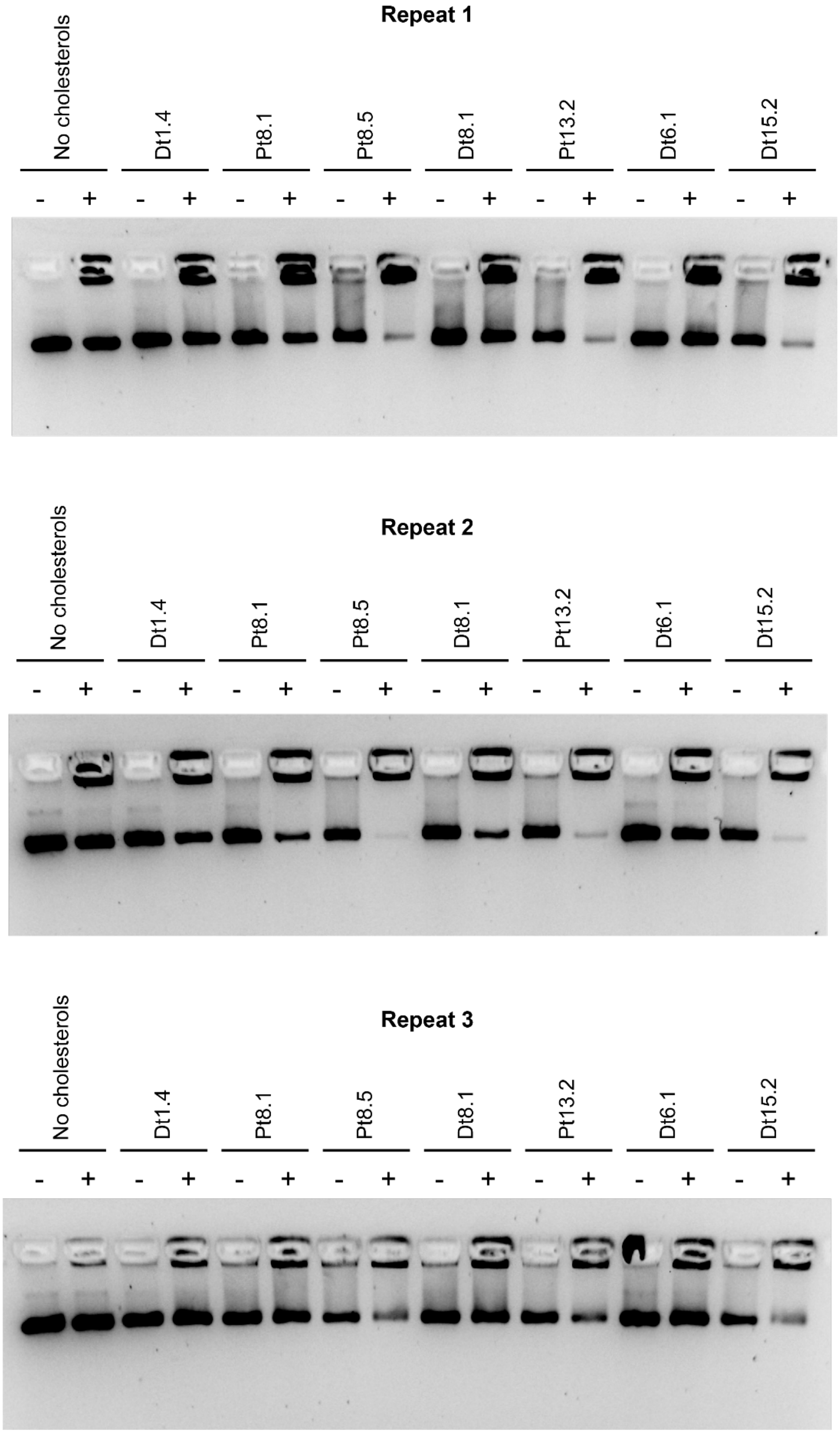


**Supplementary Figure 18 – Membrane binding vs. cholesterol linker, repeat gels**

Gel-shift assay for membrane binding with different designs of cholesterols on the tile. Three repeats were performed.

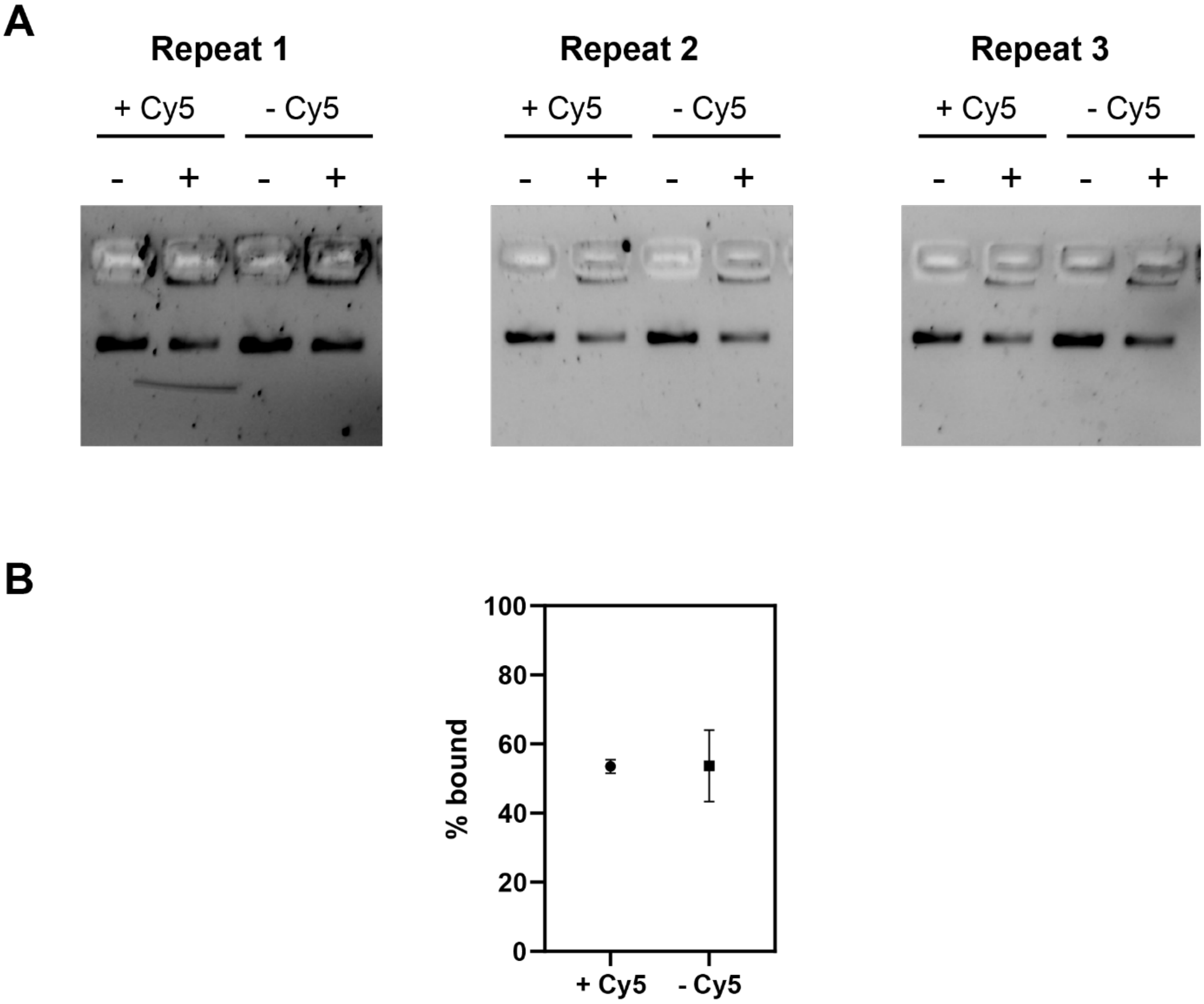


**Supplementary Figure 19 – Membrane binding of tile with/without cholesterols**

A. Gel-shift assay for membrane binding of the tile (Pt8.1) with and without Cy5 decorated on the tile. The location of staple extension for Cy5 decoration is given in Supplementary Figure 8. The gels were imaged in the SyBr Safe channel. B. Tabulated results from the gels in A.

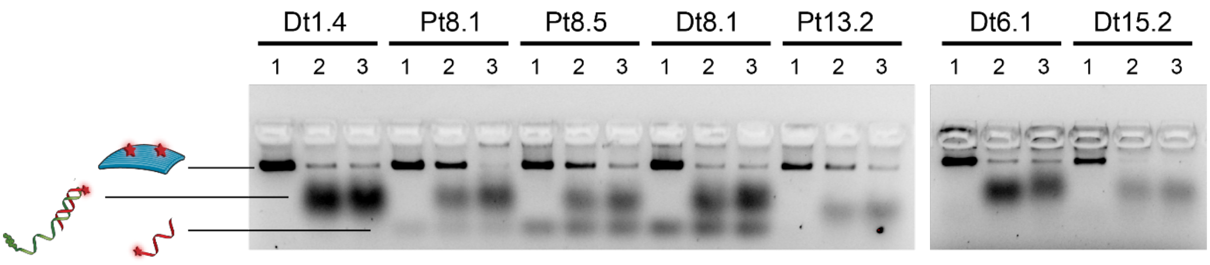


**1** No displacement

**2** Displacement at room temperature

**3** Displacement at 37C

**Supplementary Figure 20 – Strand displacement of cholesterol from tile**

Gel image showing the strand displacement of the cholesterol strands off the tile. The quantification of this gel is given in Figure 6A.iii.

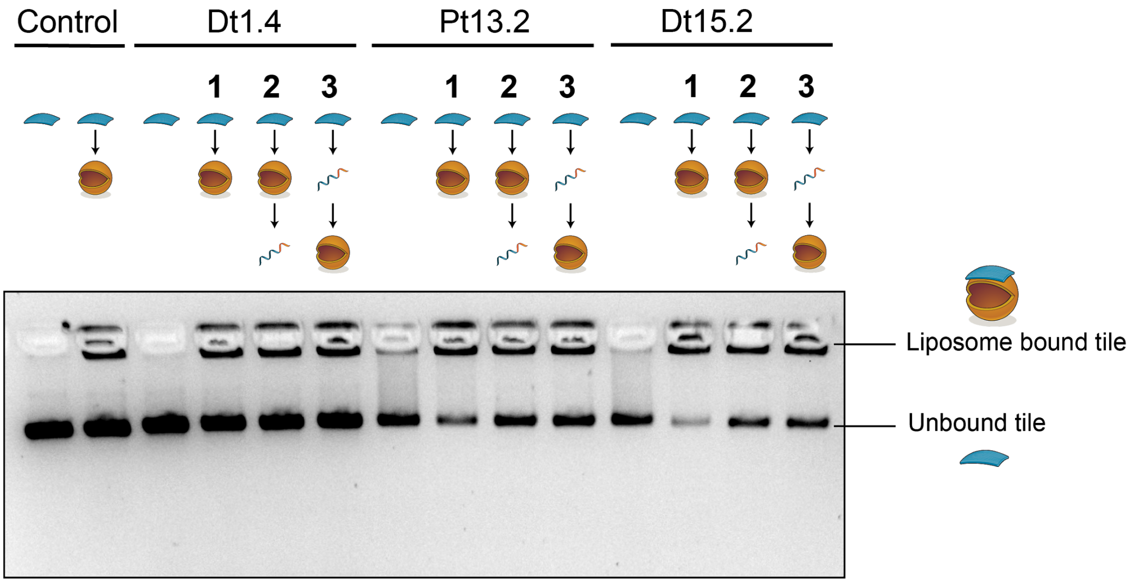


**Supplementary Figure 21 – Membrane binding and displacement of tiles**

Gel showing binding and unbinding of the tile from the SUVs. In this experiment, the fluorescent strand attachment onto the tile is independent of the cholesterol attachment to the tile. For each design, three different conditions were tested: (1) tile + extruded liposomes, (2) tile + extruded liposomes + displacer and (3) pre-displaced tile + extruded liposomes. The control is a tile with no cholesterols on it (lane 1). The results from these gels were tabulated and are shown in Figure 6B.

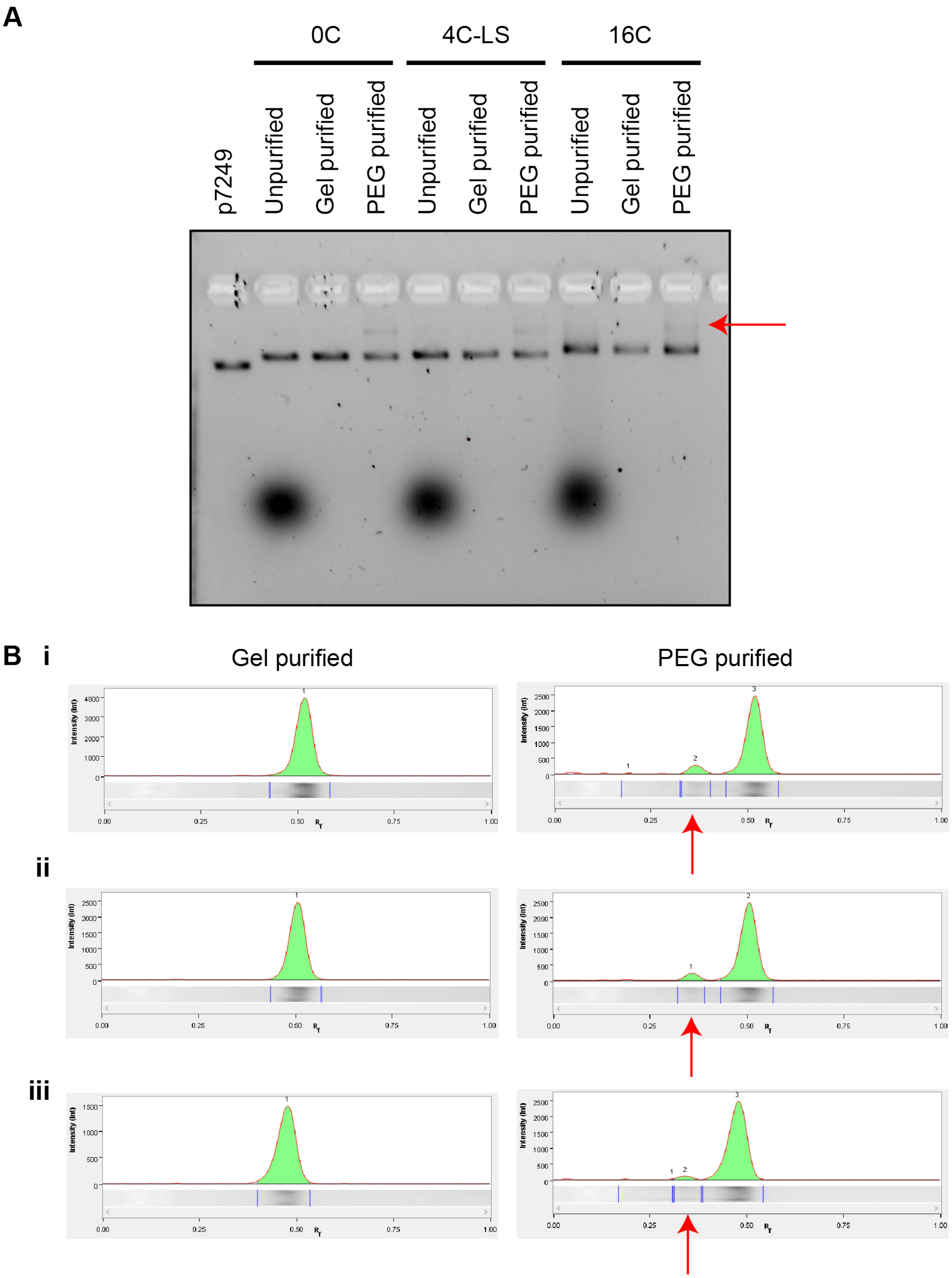


**Supplementary Figure 22 – Comparison of gel and PEG purification**

Effect of gel purification and PEG purification. A. Gel image comparing the different purification methods. B. Integrated volume of the gel bands showing aggregation (red arrow points to the aggregation of the tiles. i. 0C. ii. 4C. iii. 16C.

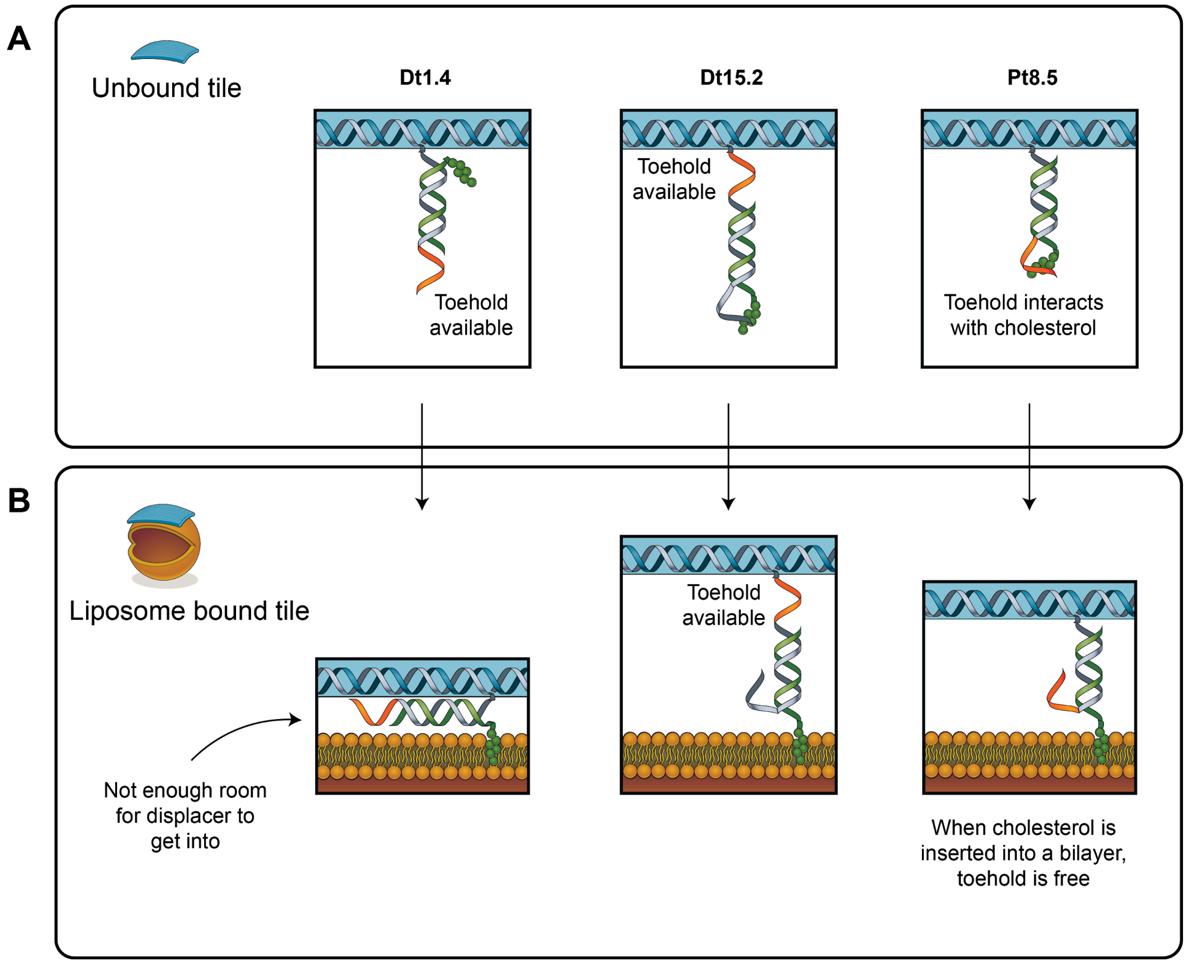


**Supplementary Figure 23– Schematic of interactions between toehold and cholesterol**

Hypothesised behaviours of toehold when the tile is unbound (A) and when the tile is bound to liposomes (B) for Dt1.4, Dt15.2 and Pt8.5. In Dt1.4, the toehold which is distal to the cholesterol is available when the tile is not bound. However, when the tile is bound to an extruded liposome, the small spacing between the tile and liposomes may prevent the displacement strand from getting to the toehold. In Dt15.2, the toehold is available both when the tile is bound and unbound to the liposomes. In Pt8.5, the toehold is expected to interact with cholesterol in the unbound state, which explains the lack of displacement at room temperature shown in Figure 6A.iii. However, when the cholesterol is inserted into the bilayer, it is hypothesised that the interaction between the cholesterol and the toehold is diminished, resulting in a more accessible toehold.
